# Organizing memories for generalization in complementary learning systems

**DOI:** 10.1101/2021.10.13.463791

**Authors:** Weinan Sun, Madhu Advani, Nelson Spruston, Andrew Saxe, James E. Fitzgerald

## Abstract

Memorization and generalization are complementary cognitive processes that jointly promote adaptive behavior. For example, animals should memorize a safe route to a water source and generalize to features that allow them to find new water sources, without expecting new paths to exactly resemble previous ones. Memory aids generalization by allowing the brain to extract general patterns from specific instances that were spread across time, such as when humans progressively build semantic knowledge from episodic memories. This cognitive process depends on the neural mechanisms of systems consolidation, whereby hippocampal-neocortical interactions gradually construct neocortical memory traces by consolidating hippocampal precursors. However, recent data suggest that systems consolidation only applies to a subset of hippocampal memories; why certain memories consolidate more than others remains unclear. Here we introduce a novel neural network formalization of systems consolidation that highlights an overlooked tension between neocortical memory transfer and generalization, and we resolve this tension by postulating that memories only consolidate when it aids generalization. We specifically show that unregulated memory transfer can be detrimental to generalization in unpredictable environments, whereas optimizing systems consolidation for generalization generates a high-fidelity, dual-system network supporting both memory and generalization. This theory of generalization-optimized systems consolidation produces a neural network that transfers some memory components to the neocortex and leaves others dependent on the hippocampus. It thus provides a normative principle for reconceptualizing numerous puzzling observations in the field and provides new insight into how adaptive behavior benefits from complementary learning systems specialized for memorization and generalization.

## INTRODUCTION

The brain’s ability to learn, store, and transform memories lies at the heart of our ability to make adaptive decisions^1^. Memory is threaded through cognition, from perception through spatial navigation to decision-making and explicit conscious recall. Befitting the central importance of memory, brain regions including the hippocampus appear specifically dedicated to this challenge^2–4^.

The concept of memory has refracted through psychology and neurobiology into diverse subtypes and forms that have been difficult to reconcile. Taxonomies of memory have been drawn on the basis of psychological content, for instance differences between memories for detailed episodes and semantic facts^5^; on the basis of anatomy, for instance differences between memories that are strikingly dependent on hippocampus versus those that are not^6^; and on the basis of computational properties, for instance differences between memories reliant on pattern-separated or distributed neural representations^7^. Many previous theories have tried to align and unify psychological, neurobiological, and computational memory taxonomies^8–12^. However, none has yet resolved long-standing debates on where different kinds of memories are stored in the brain, and, fundamentally, why different kinds of memories exist.

Classical views of systems consolidation, such as the standard theory of systems consolidation^8,13,14^, have held that memories reside in hippocampus before transferring completely to neocortex. Related neural network models, such as the complementary learning systems theory, have further offered a computational rationale for systems consolidation based on the benefits of coupling complementary fast and slow learning systems for integrating new information into existing knowledge^9,15–17^. However, these theories lack explanations for why some memories remain forever hippocampal dependent, as shown in a growing number of experiments^18,19^. On the other hand, more recent theories, like multiple trace theory^11,20^ and trace transformation theory^21,22^, hold that the amount of consolidation can depend on memory content, but they do not provide quantitatively clear criteria for what content will consolidate, nor why this might be beneficial for behavior.

One possible way forward is to see that memories serve not only as veridical records of experience, but also to support generalization in novel circumstances^23–29^. For instance, individual memorized experiences almost never repeat exactly, but they allow us to identify systematic relationships between features of the world, such as “ravines predict the presence of water,” which are common and important for behavior. Here we introduce a mathematical neural network theory of systems consolidation founded on the principle that memory systems and their interactions collectively optimize generalization. Our theory mathematically defines the generalization performance of an algorithm as its expected error for any possible input, including new ones. This definition is widespread in statistics and machine learning, and it resonates with the intuitive notion that generalizations apply regularities inferred from specific instances to new circumstances. The resulting theory offers new perspectives on diverse experimental phenomena and explains why interaction between multiple brain areas is beneficial. Accurate generalizations require consistent relationships within the environment, and our theory optimizes generalization by using the predictability of a memorized experience to determine when and where its memory trace resides. Our results thus propose a quantitative and unified theory of the organization of memories based on their utility for future adaptive behavior.

## RESULTS

### Formalizing systems consolidation

We conceptualize an animal’s experiences in the environment as structured neuronal activity patterns that the hippocampus rapidly encodes and the neocortex gradually learns to produce internally^9,17,30,31^ (Fig. 1a). We hypothesize that systems consolidation allows neocortical circuits to learn many structured relationships between different subsets of these active neurons. Focusing on one of these relationships at a time, neocortical circuitry might learn through many experiences (Fig. 1b) to produce the responses of a particular *output* neuron from the responses of other *input* neurons (Fig. 1c). For example, in a human, an output neuron contributing to a representation of the word “bird” might receive strong inputs from neurons associated with wings and flight. In a mouse, an output neuron associated with behavioral freezing might receive strong inputs from neurons associated with the sound of an owl, the smell of a snake, or the features of a laboratory cage where it had been shocked^32^.

**Fig. 1.**
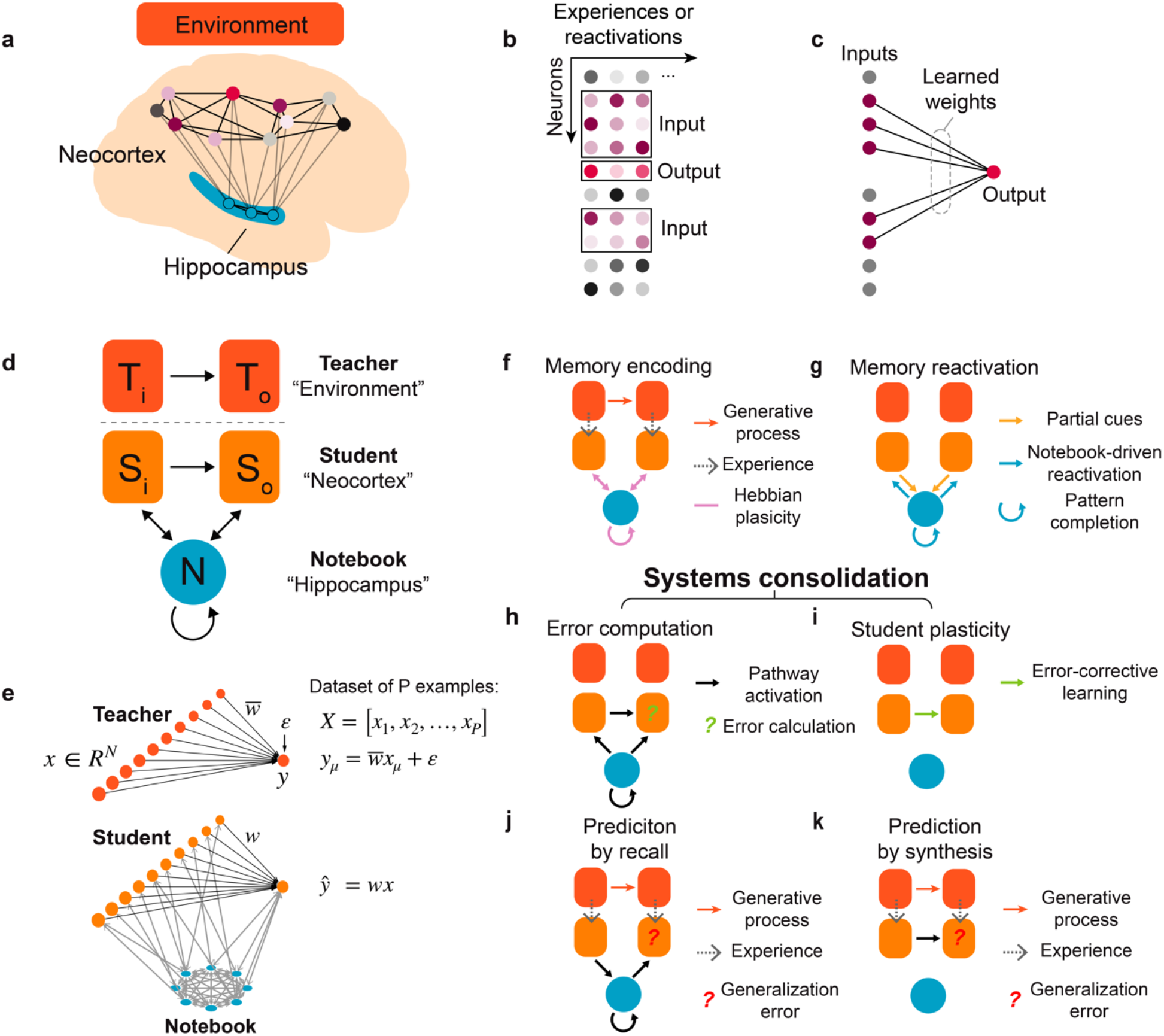
Neural network model of systems consolidation. **(a-c)** Our theoretical framework assumes that neocortex extracts and encodes environmental relationships within the weights between distributed neocortical neurons in a process mediated by hippocampal reactivation. We color the output neuron in red, and its activity is determined by the purple input neurons that are connected to it. This example is for illustrative purposes, and an input neuron in one relationship could be an output neuron in another relationship. **(d)** Cartoon of the teacher-student-notebook formalism; subscripts “i” and “o” refer to input and output layers. **(e)** Neural network model architecture used in most simulations, unless otherwise noted. The teacher is a linear, shallow network with fixed weights that transforms N dimensional input into a scalar y, with a noise term ε added to vary the signal-to-noise ratio of the teacher. The student is typically a size-matched network to the teacher, with trainable weights *w.* The notebook is a Hopfield network that is bidirectionally connected to the student which serves as a one-shot learning module for memory encoding and replay (see Methods for details). **(f-k)** Stages of learning and inferences in the model. Student is activated by each of the teacher-generated example while notebook encode this example through one-shot Hebbian plasticity (**f**). The notebook can reactivate the encoded examples offline and reactivate the student (**g**). Notebook could reactivate previously encoded memories offline to induce memory recall in the student (**h**) and student learning (**i**). The student can use either the notebook or the student for inference (**j, k**).

We first sought to develop a theoretically rigorous mathematical framework to formalize this view of how systems consolidation contributes to learning. Our framework builds on the complementary learning systems hypothesis^9,17^, which posits that fast learning in hippocampus guides slow learning in neocortex to provide an integrated learning system that outperforms either subsystem on its own. Here we formalize this notion as a neocortical *student* that learns to predict an environmental *teacher,* aided by past experiences recorded in a hippocampal *notebook* (Fig. 1d). Note that although the theory is centered around hippocampal-neocortical interactions, the core theoretical principles can be potentially applied to other brain circuits that balance fast and slow learning^33–35^.

We modeled each of these theoretical elements with a simple neural network amenable to mathematical analyses (Fig. 1e, Methods). Specifically, we modeled the teacher as a linear feedforward network that generates input-output pairs through fixed weights with additive output noise, the student as a size-matched linear feedforward network with learnable weights^36–38^, and the notebook as a sparse Hopfield network^39–41^. The student learns its weights from a finite set of examples (experiences) that contain both signal and noise. We modeled the standard theory of systems consolidation by optimizing weights for memory. This means that the squared difference between the teacher’s output and the student’s prediction should be as small as possible, averaged across the set of past experiences. Alternatively, we hypothesize that a major goal of the neocortex is to optimize generalization. This means that the squared difference between the teacher’s output and the student’s prediction should be as small as possible, averaged across possible future experiences that could be generated by the teacher.

Learning starts when the teacher activates student neurons (Fig. 1f, gray arrows). The notebook encodes this student activity by associating it with a random pattern of sparse notebook activity using Hebbian plasticity (Methods; Fig. 1f, pink arrows). This effectively models hippocampal activity as a pattern-separated code for indexing memories^42,43^. The recurrent dynamics of the notebook network implement pattern completion^39,44^, whereby full notebook indices can be reactivated randomly from spontaneous activity or purposefully from partial cues^45^ (Methods; Fig. 1g). Student-to-notebook connections allow the student to provide the partial cues that drive pattern completion (Fig. 1g, orange arrows). Notebook-to-student connections then allow the completed notebook index to reactivate whatever student representations were active during encoding (Fig. 1g, blue arrows). Taken together, these three processes permit the student to use the notebook to recall memories from related experiences in the environment. Thus, our theory concretely models how the neocortex could use the hippocampus for memory recall.

We model systems consolidation as plasticity of the student’s internal synapses (Figs. 1h, 1i). The student’s plasticity mechanism is guided by notebook reactivations (Fig. 1h), similar to how hippocampal replay is hypothesized to contribute to systems consolidation^46^. Slow, error-corrective learning aids generalization^47^, and here we adjust internal student weights with gradient-descent learning (Fig. 1i). Specifically, we assume that offline notebook reactivations provide targets for student learning (Methods), where the notebook-reactivated student output is compared with student’s internal prediction to calculate an error signal for learning. We consider models that set the number of notebook reactivations to optimize either memory transfer or generalization. The integrated system can use the notebook (Fig. 1j) or only the learned internal student weights (Fig. 1k) to make output predictions from any input generated by the teacher. We will show that each pathway has distinct advantages for memory and generalization.

### Generalization-optimized complementary learning systems (Go-CLS)

We next simulated the dynamics of memorization and generalization in the teacher-student-notebook framework to investigate the impact of systems consolidation. We first modeled the standard theory of systems consolidation as limitless notebook reactivations that optimized student memory recall (Fig. 2a, c, e, Methods). Learning begins when the notebook stores a small batch of examples, which are then repetitively reactivated by the notebook in each epoch to drive student learning (Methods). In separate simulations, examples were generated by one of three teachers that differed in their degree of predictability, here controlled by the signal-to-noise ratio (SNR) of the teacher network’s output (Fig. 1e, Methods). The notebook was able to accurately recall the examples provided by each teacher from the beginning (Fig. 2a, c, e, *dashed blue lines*), and we showed mathematically that recall accuracy scaled with the size of the notebook (Supplementary Material 5.2). Notebook-mediated generalization (Student In → Notebook → Student Out) was poor for all three teachers (Fig. 2a, c, e, *dashed red lines*), as rote memorization poorly predicts high-dimensional stimuli that were not previously presented or memorized (Supplementary Material 5.3). The student gradually reproduced past examples accurately (Fig. 2a, c, e, *solid blue lines*), but the signal in each example was contaminated by whatever noise was present during encoding and repetitively replayed throughout learning. Therefore, although the generalization error decreased monotonically for the noiseless teacher (Fig. 2a, *solid red line*), noisy teachers resulted in the student eventually generalizing poorly (Fig. 2c, e, *solid red lines*). From a mathematical point of view, this is expected, as the phenomenon of overfitting to noisy data is well appreciated in statistics and machine learning^48,49^.

**Fig. 2.**
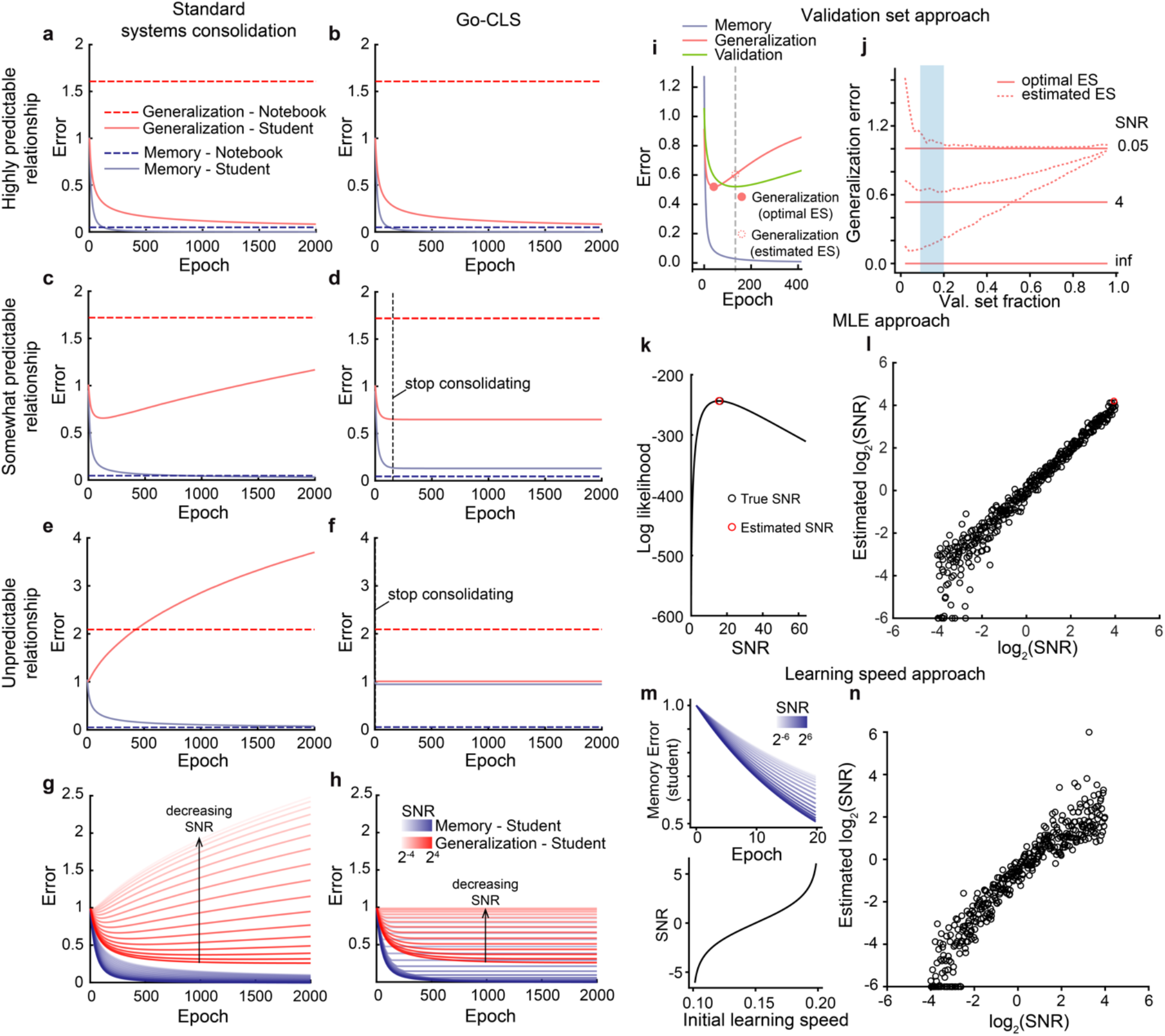
The predictability of experience controls the dynamics of systems consolidation. **(a-h)** Dynamics of student generalization error, student memorization error, notebook generalization error, and notebook memorization error when optimizing for student memorization (a, c, e, g) or generalization (b, d, f, h) performance. The student’s input dimension *N* = 100, and the number of patterns stored in the notebook *P* = 100 (all encoded at epoch = 1; epochs in the x-axis correspond to the time passage during systems consolidation). Notebook contains *M* = 2000 units, with a sparsity *a* = 0.05. During each epoch, 100 patterns are randomly sampled from the *P* stored patterns for reactivation and training the student. The student’s learning rate is 0.015. Teachers differed in their levels of predictability (a, b: SNR = ∞; c, d: SNR = 4; e, f: SNR = 0.05; g, h: SNR ranges from 2^-4^ to 2^4^). (**i-n**) Mechanistic methods for estimating predictability. **(i)** Using a validation set to estimate optimal early stopping time (SNR = 4, P = N = 100, 10% of P are used as validation set and not used for training). Filled red dot marks the generalization error at the optimal early stopping time (optimal ES); Dashed red dot marks the generalization error at the early stopping time estimated by the validation set (estimated ES). **(j)** Generalization error at optimal (solid red line) vs estimated early stopping time (dashed red line), as a function of validation set fraction, SNR, and alpha (P/N). The blue shading indicates validation set fraction from 10% to 20%. **(k)** Illustration of maximum likelihood estimation (MLE) (Supplementary Material 9.2). **(l)** MLE predicts true SNR well from teacher-generated data. **(m)** initial learning speed monotonically increases as a function of SNR. (**n**) Initial learning speed serves as a good feature for estimating true SNR in numerical simulations (P = N = 1000).

The implications of these findings for psychology and neuroscience are far reaching, as the standard theory of systems consolidation assumes that generalization follows naturally from hippocampal memorization and replay; it does not consider when systems consolidation is detrimental to generalization. For example, previous neural network models of complementary learning systems focused on learning scenarios where the mapping from input to output was fully reliable^8,9^. Within our teacher-student-notebook framework, this means that the teacher is noiseless and perfectly predictable by the student architecture. In such scenarios, standard systems consolidation continually improved both memorization and generalization in our model (Fig. 2a, *solid red line*). However, for less predictable environments, our theory suggests that too much systems consolidation can severely degrade generalization performance by leading the neocortex to overfit to unpredictable elements of the environment (Fig. 2c, *solid red).* In highly unpredictable environments, any systems consolidation at all can be detrimental to generalization (Fig. 2e, *solid red*). If the goal of systems consolidation is full memory transfer, then our theory illustrates that the system pays a price in reduced ability to generalize in uncertain environments.

Given the intricate effect of training example replay on generalization, what systems consolidation strategy would optimize generalization? Here we propose a new theory— generalization-optimized complementary learning systems (Go-CLS)—which considers the normative hypothesis that the amount of systems consolidation is adaptively regulated to optimize the student’s generalization accuracy based on the predictability of the input-output mapping (Fig. 2b, d, f). For the teacher with a high degree of predictability, the student’s generalization error always decreased with more systems consolidation (Fig. 2b, *solid red line),* and the student could eventually recall all stored memories (Fig. 2b, *solid blue line*). Memory transfer therefore arises as a property of a student that learns to generalize well from this teacher’s examples. In contrast, a finite amount of consolidation (here modeled by a fixed number of notebook reactivations) was necessary to minimize the generalization error when the teacher had limited predictability (Fig. 2d, f), and our normative hypothesis is that systems consolidation halts at the point where further consolidation harms generalization (Fig. 2d, f, *vertical black dashed line*). The resulting student could generalize nearly optimally from each of the teachers’ examples (Fig. 2d, f, *solid red lines,* Supplementary Material 7.2), but its memory performance was hurt by incomplete memorization of the training data (Fig. 2d, f, *solid blue lines*). Nevertheless, the notebook could still recall the memorized examples (Fig. 2b, d, f, *dashed blue lines*). Go-CLS thus results in an integrated system that can both generalize and memorize by using two systems with complementary properties.

These examples show that the dynamics of systems consolidation models depend on the degree of predictability of the teacher. We therefore leveraged our analytical results to comprehensively compare the standard theory of systems consolidation to Go-CLS theory for all degrees of predictability (Supplementary Material 6, 7). Standard systems consolidation eventually consolidated all memories for any teacher (Fig. 2g, blue). As anticipated by Fig. 2a-f, the generalization performance varied dramatically with the teacher’s degree of predictability (Fig. 2g, red). Generalization errors were higher for less predictable teachers, and optimal consolidation amounts were lower. Therefore, Go-CLS removed the detrimental effects of overfitting (Fig. 2h, red) but ended before the student could achieve perfect memorization (Fig. 2h, blue, non-zero error). Both the generalization performance and the memory performance improved as the teacher’s degree of predictability increased (Fig. 2h).

Fully implementing this strategy for Go-CLS requires a supervisory process capable of estimating the optimal amount of consolidation (Supplemental Material 9). One conceptually simple way to do this is to directly estimate the generalization error dynamics (Fig. 2i-j), which would not require explicit inference of the teacher’s predictability. For instance, the brain could divide the notebook’s memorized examples into a training set that drives student learning and a validation set that does not. Since the student’s error on the validation set is an estimate of the generalization error, the brain could regulate consolidation by stopping student learning when the validation error starts increasing (Fig. 2i). This strategy works best for relatively small validation sets, as this permits learning from most examples (Fig. 2j).

Another strategy to regulate consolidation is to estimate the predictability of the teacher (Fig. 2k-n). Although the computational algorithms and biological mechanisms are unknown, we know that humans estimate the predictability of information routinely. For example, when a little-league pitcher takes the mound, we may estimate the likelihood that they will throw a strike. In the teacher-student-notebook model, the system could statistically estimate the teacher’s degree of predictability as the one that maximizes the likelihood of the teacher-generated examples (Fig. 2k). This amounts to comparing the input-output covariance of the teacher-generated data to theoretical expectations, which varies in a predictable way with SNR (Supplementary Material 9). Alternatively, the brain could use the simpler heuristic that the initial learning speed (for a given sized dataset) correlates with predictability (Fig. 2m, Supplementary Material 9). Each of these methods provides a reasonably accurate estimate of the teacher’s degree of predictability (Fig. 2l, n), which could be used to estimate the optimal early stopping time (see Fig. 2g). Such estimates rely on prior knowledge relating data statistics to the teacher’s degree of predictability, which for more complex environments could be established by meta-learning over developmental, lifelong, and evolutionary timescales^50^.

### Relating Go-CLS to diverse experimental results

Experimental literature on the time course of systems consolidation and time-dependent generalization provides important constraints on our theory. We thus sought to model these effects by translating mean square errors (Fig. 2g, h) into memory or generalization scores, where 0 indicates random performance and 1 indicates perfect performance (Fig. 3a-d, Methods). Our framework can use either the student or the notebook to recall memories or generalize (Fig. 1j, k). Here we model the combined system by making predictions with whichever subsystem is more accurate (Methods). This assumption isn’t critical, as the combined memory (Fig. 3a, b) and generalization scores (Fig. 3c, d) often map onto the notebook and student performances, respectively, but this assumption allows the combined system to switch between subsystems over time (Supplemental Material 6.1). Other models might implement more complex memory system selection policies or combine both pathways to obtain statistically better predictions throughout learning. We simulated hippocampal lesions by preventing the combined system from using notebook outputs and ending systems consolidation at the time of the lesion (Fig. 3a, b, cyan). As it takes time for the student to learn accurate generalizations, our systems consolidation models exhibit time-dependent generalization (Fig. 3c, d, purple). In contrast, the notebook permitted accurate memory retrieval from the start (Fig. 3a, b, black).

**Fig. 3.**
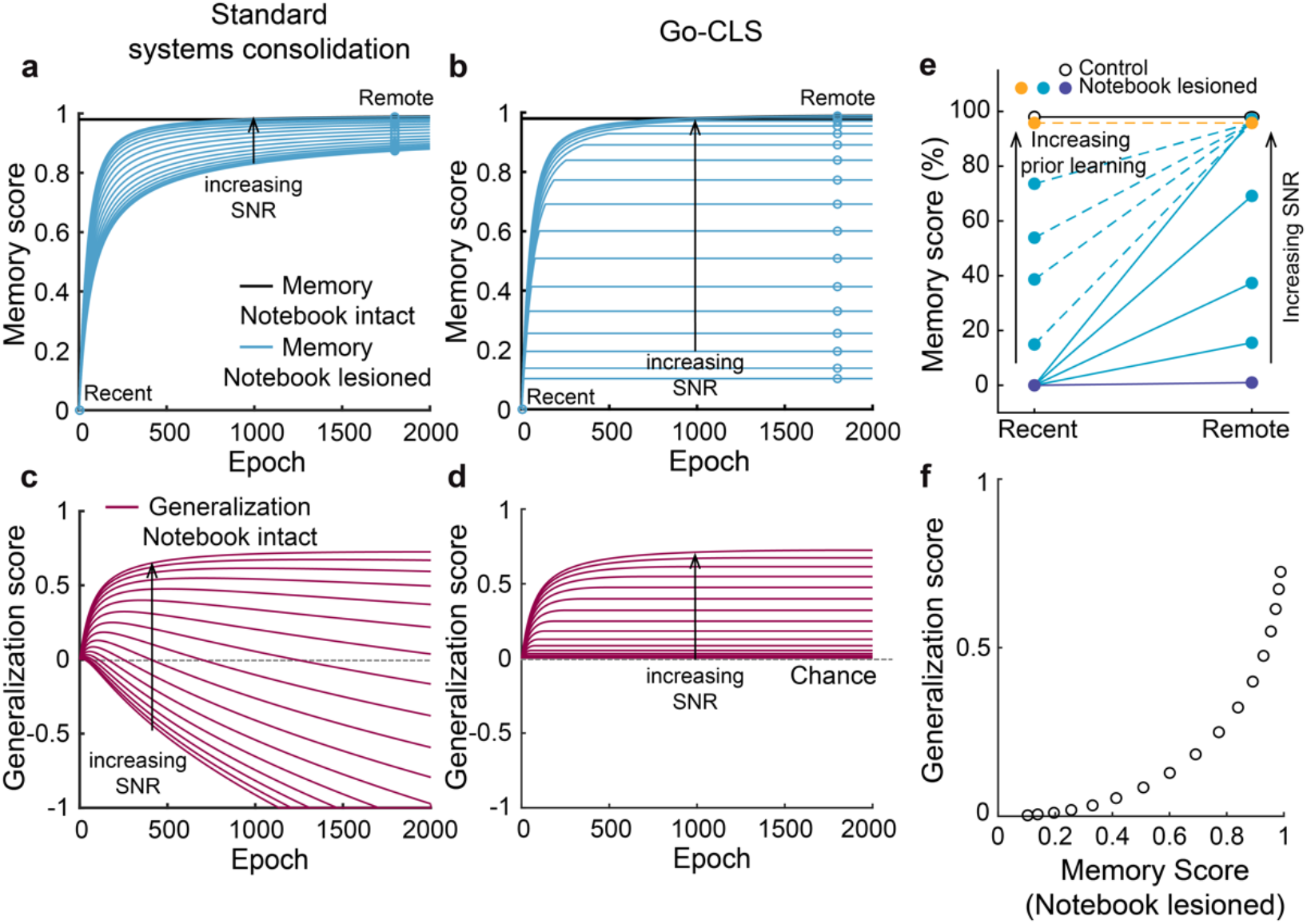
Go-CLS mirrors memory research findings. **(a,b,c,d)** Memorization (a, b) and generalization (c, d) scores for the integrated student-notebook system as a function of time and SNR, when optimized for student memorization (a, c) or generalization (b, d). Memory and generalization scores are translated from respective error values by Score = (*E*_0_ - *E_t_*)/*E_0_, E_0_* and *E_t_* are the generalization or memory errors of a zeroweight student and a trained student at epoch = *t,* respectively. The effect of notebook lesion on memory performance (cyan lines and open circles, open circles simply demarcate the cyan lines at hypothetical “recent” and “remote” timepoints) depended on optimization objective and time (a, b). **(e)** Go-CLS can reproduce the diversity of retrograde amnesia curves (see Methods for model details). **(f)** Memory and generalization scores are positively correlated after notebook lesioning.

Standard systems consolidation and Go-CLS theory make strikingly different predictions for how retrograde amnesia depends on the teacher’s degree of predictability (Fig. 3a-d). Researchers usually classify hippocampal amnesia dynamics according to whether memory deficits are similar for recent and remote memories (flat retrograde amnesia), more pronounced for recent memories (graded retrograde amnesia), or absent for both recent and remote memories (no retrograde amnesia) (Fig. 3e). As expected, notebook lesions always produced temporally graded retrograde amnesia curves in the standard theory^13^ (Fig. 3a). When systems consolidation was instead optimized for generalization, the effects of notebook lesions depended strongly on the predictability of the teacher (Fig. 3b). Therefore, Go-CLS theory can recapitulate a wide diversity of retrograde amnesia curves (Fig. 3e). High and low predictability experiences lead to graded and flat retrograde amnesia, respectively (Figs. 3a, 3e, *solid lines).* A period of prior consolidation of highly predictable experiences decreases the slope of graded retrograde amnesia (Fig. 3e, *dashed light-blue lines*), and it’s possible to see no retrograde amnesia at all when the prior consolidation was extensive (Fig. 3e, *dashed orange line,* Methods). This conceptually resembles schema-consistent learning^51^.

Experiments on time-dependent generalization can also differentiate between Go-CLS theory and the standard theory. Diverse generalization curves resulted from either model of systems consolidation (Fig. 3c, d), with maximal generalization performance increasing with the predictability of the teacher. However, student overfitting meant that only Go-CLS maintained this performance over time. Standard systems consolidation could even result in a student generalizing maladaptively, resulting in worse-than-chance performance where the trained student interpolates noise in past examples to produce wildly inaccurate outputs (Fig. 2c). Most fundamentally, Go-CLS theory predicts that memory transfer and generalization improvement should be correlated with each other (Fig. 3f), as systems consolidation leads to both. Unpredictable experiences should not consolidate because this would cause maladaptive generalization. Such memories are thus left in their original form and susceptible to strong retrograde amnesia following hippocampal lesion. In contrast, predictable experiences should consolidate and be associated with weak retrograde amnesia and useful learned generalizations.

These results may allow Go-CLS theory to reconcile a wide diversity of experimental findings (Supplementary Material 11). Go-CLS potentially resolves apparent conflicts in the literature as arising from differing degrees of predictability in the underlying experimental paradigms. This hypothesized correspondence between past experiments and their predictability is intriguing but inconclusive, as it is not yet clear how to quantify the degree of predictability for arbitrary experiments and real-world scenarios. In other words, the theory is consistent with existing findings in principle, but its post-diction of them requires plausible assumptions that may be wrong. Future experiments are therefore critical (see detailed discussions of experimental tests in Supplementary Material 12). Our core theoretical prediction is that the brain evolved to optimize for generalization by regulating the amount of systems consolidation based on the predictability of experience^52^. Direct tests of this prediction require experimental task designs that intentionally vary the degree of predictability and assess the effect on systems consolidation. In addition, experiments that identify the biological mechanisms of predictability estimation and consolidation regulation would be required to establish a comprehensive picture of the neural correspondence of the Go-CLS theory.

### Normative benefits of complementary learning systems for generalization

Our framework also provides theoretical insights into the complementary learning systems hypothesis, which posits that hippocampal and neocortical systems exploit fundamental advantages provided by coupled fast and slow learning modules^9,17,53–55^. We first investigated its basic premise by comparing generalization in the optimally regulated student-notebook network (Fig. 4a) to what is achievable with isolated student (Fig. 4b) and notebook networks (Fig. 4c). Since the student models the neocortex and the notebook models the hippocampus, these isolated student and notebook networks model learning with only neocortex or only hippocampus, respectively.

**Fig. 4.**
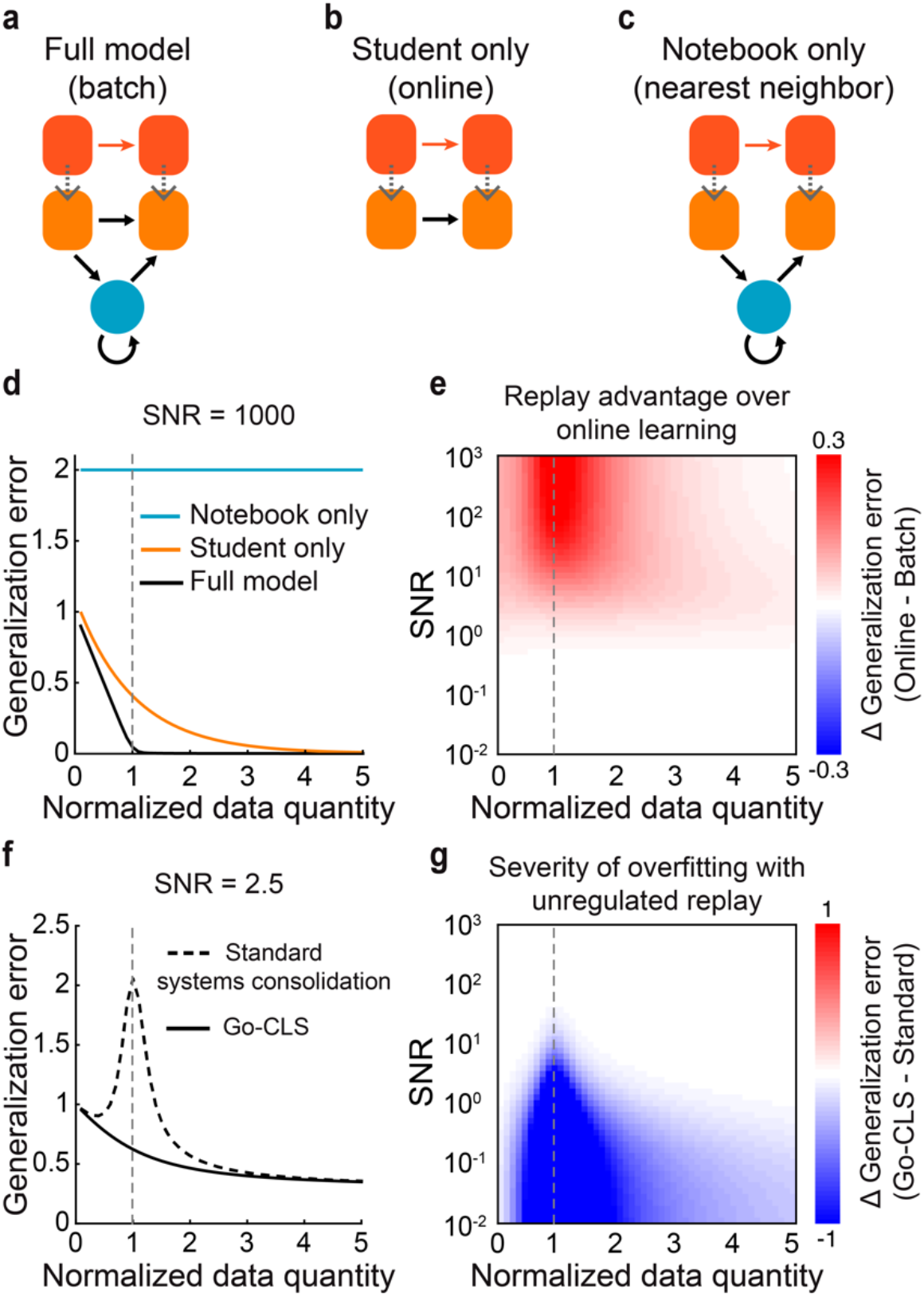
Normative benefits of complementary learning systems for generalization. **(a-c)** Schematics illustrating learning systems that can use both the student and the notebook (a), only the student’s weights (b), and only the notebook weights (c) for inference. In machine learning terminology, these systems implement batch learning, online learning, and nearest neighbor regression. **(d)** Generalization error as a function of normalized data quantity (or *alpha (a),* defined as *a* = *P/N)* for each learning system (SNR = 1000), dashed gray line indicates *a* = 1. **(e)** Advantage of Go-CLS over optimal online learning as a function of SNR and normalized data quantity, measured by the difference in generalization error. **(f)** Generalization error as a function of normalized data quantity for the combined system, learning either through Go-CLS or standard systems consolidation (SNR = 2.5). **(g)** Severity of overfitting, measured by the difference in generalization error between standard systems consolidation and Go-CLS.

Both the degree of predictability and the amount of available data impact the time course of systems consolidation in the student-notebook network (Supplementary Material 6, 7), so we used our analytical solutions to systematically examine how late-time memory and generalization jointly depend on the amount of training data and degree of predictability (Supplementary Fig. 2). With just a student (Fig. 4b), the system must learn online from each example with no ability to revisit it. This limitation prevented the optimal student-only network from generalizing as efficiently from predictable teacher-generated data compared to the optimal student-notebook network (Fig. 4d, *yellow vs black curves*), despite modulating its learning rate online to achieve best-case generalization performance (Supplementary Material 4.2). We also confirmed that both student-containing networks generalized better than the notebook-only network (Fig. 4d). This is expected, because in high dimensions any new random pattern is almost always far from the nearest memorized pattern (Supplementary Material 5.3); this is the so-called “curse of dimensionality”^56^.

The generalization gain provided by the student-notebook network over the student-only network was most substantial when the teacher provided a moderate amount of predictable data (Figs. 4d-e, *dashed gray*). This result follows because the student-notebook network was unable to learn much when the data were too few or too noisy, and notebook-driven encoding and reactivation of data was unnecessary when the student had direct access to a large amount of teacher-generated data (Supplementary Material 4, 7). Hence an integrated dual memory system was normatively superior when experience was available, but limited, and the environment was at least somewhat predictable.

The ability to replay examples was most advantageous when the number of memorized examples equaled the number of learnable weights in the student (Fig. 4e, *dashed gray*). Remarkably, this amount of data was also the worst-case scenario for overfitting to noise in standard systems consolidation (Figs. 4f-g, *dashed gray,* Supplementary Fig. 2c), similar to the “double descent” phenomenon in machine learning^37,57^, where the worst overfitting also happens at an intermediate amount of data related to the network size. Intuitively, neural networks must tune their weights most finely when the number of memorized patterns is close to the maximal achievable number (capacity). This often requires drastic changes in weights to reduce a small training error residual, producing noise-corrupted weights that generalize poorly. The optimal student-notebook network avoided this issue by regulating the amount of systems consolidation according to the predictability of the teacher. We propose that the brain might similarly regulate the amount of systems consolidation according to the predictability of experiences (see Discussion).

### Many facets of unpredictability

Our simulations and analytical results show that the degree of predictability controls the consolidation dynamics that optimize generalization. We emphasized the example of a linear student (Fig. 5a) that learns from a noisy linear teacher (Fig. 5b). However, inherent noise is only one of several forms of unpredictability that can cause poor generalization without regulated systems consolidation. For example, when the teacher implements a deterministic transformation that is impossible for the student architecture to implement, the unmodellable parts of the teacher mapping are unpredictable and act like noise (Supplementary Material 10). For instance, a linear student cannot perfectly model a nonlinear teacher (Fig. 5c). Similarly, when the teacher’s mapping involves relevant input features that the student cannot observe, the contribution of the unobserved inputs to the output is generally impossible to model (Fig. 5d). This again results in unpredictability from the student’s perspective. These sources of unpredictability all consist of a modellable signal and an unmodellable residual (noise) (Supplementary Material 10), and they yield similar training and generalization dynamics (Fig. 5e, f). The real world is noisy and complicated, and the brain’s perceptual access to relevant information is limited. Realistic experiences thus frequently combine these sources of unpredictability.

**Fig. 5.**
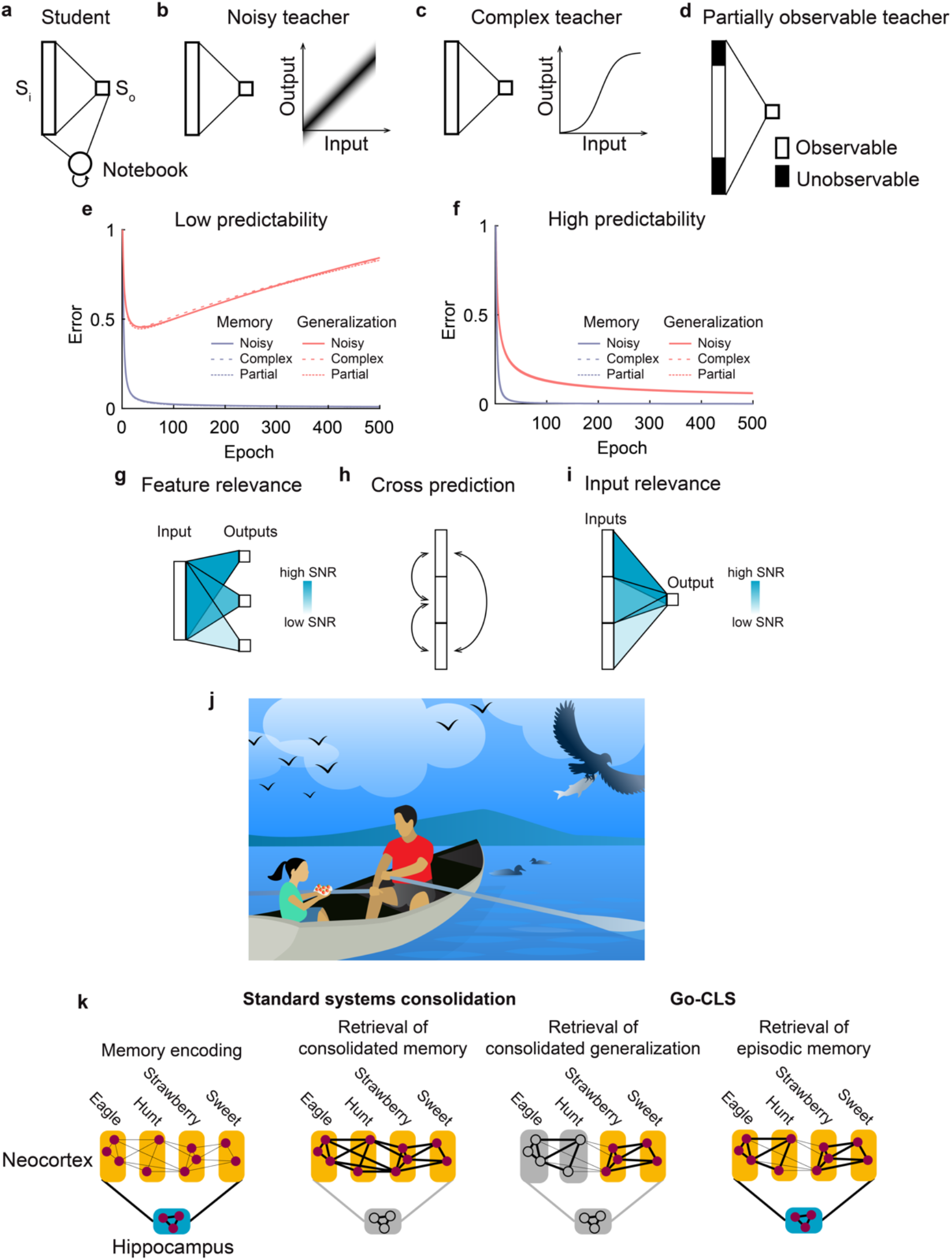
Many forms of unpredictability demand regulated systems consolidation. **(a)** The student-notebook learning system. **(b-d)** Example teachers with unpredictable elements. **(b)** A teacher that linearly transforms inputs into noisy outputs. (**c**) A teacher that applies a nonlinear activation function at the output unit. **(d)** A teacher that only partially reveals the relevant inputs to the learning system. **(e, f)** Varying predictability within the three different teachers all lead to quantitatively similar learning dynamics (complex teacher implements a sine function at the output unit, see Methods for simulation details). **(g-i)** The degree of predictability can vary in many ways. For example, the same inputs can differentially predict various outputs (g), features can cross predict each other with varying levels of predictability (h), and different learning systems could attend to different teacher features to predict the same output (i). (**j**) Cartoon illustrating a child’s experience at a lake with her father. **(k)** A cartoon illustrating conceptual differences between what is consolidated in standard systems consolidation and Go-CLS.

All the above-mentioned cases can be generally understood within the framework of approximation theory^58^. The unmodellable part represents a nonzero optimal approximation error for the student-teacher pair. For this unmodellable part to be generalization limiting, the student must also be expressive enough that the student weights overfit when attempting to fit limited data perfectly. Overfitting is also seen in more complex model architectures (Supplementary Material 10.2), such as modern deep learning models^57,59^ and we expect that the essential concepts presented here will also apply to broader model classes. For all these types of generalization-limiting unpredictability, generalization is optimized when systems consolidation is limited for unpredictable experiences. Importantly, not all unpredictability limits generalization (Supplementary Material 8). For example, independent noise during inference can actually promote generalization, such as in dropout regularization^60^.

Previously we have focused on the scenario of learning a single mapping. All real-life experiences are composed of many components, with relationships that can differ in predictability. Many relationships therefore must be learned simultaneously, and these representations are widely distributed across the brain. For instance, the same input features may have different utility in predicting several outputs (Fig. 5g). Furthermore, neocortical circuits may cross-predict between different sets of inputs and outputs (Figs. 1e, 5h); for example, perhaps predicting auditory representations from visual representations and vice versa. In this setting, each cross prediction has its own predictability determined by the noise, the complexity of the mapping, and the features it is based upon. Predictability may also depend on overt and/or covert attention processes in the student. For example, a student may selectively attend to a subset of the inputs it receives (Fig. 5i), making the predictability of the same external experience dependent on internal states that can differ across individuals, which might partially underlie the individual variability in memory consolidation seen in animal behavior^61^. For all the above-mentioned scenarios, Go-CLS theory requires the student to optimize generalization by regulating systems consolidation according to the specific degree of predictability of each modelled relationship contained in an experience. The theory therefore provides a novel predictive framework for quantitatively understanding how diverse relationships within memorized experiences should differentially consolidate to produce optimal general-purpose neocortical representations.

## DISCUSSION

The theory presented here — Go-CLS — provides a normative and quantitative framework for assessing the conditions under which systems consolidation is advantageous or deleterious. As such, it differs from previous theories that sought to explain experimental results without explicitly considering when systems consolidation could be counterproductive^8,9,13,19–21^. The central claim of this work is that systems consolidation from the hippocampus to neocortex is most adaptive if it is regulated such that it improves generalization, an essential ability enabling animals to make predictions that guide behaviors promoting survival in an uncertain world. Crucially, we show that unregulated systems consolidation results in inaccurate predictions by neural networks when limited data contain a mixture of predictable and unpredictable components. These errors result directly from the well-known overfitting problem that occurs in artificial neural networks when weights are fine-tuned to account for data containing noise and/or unlearnable structure^37,38,48,49,57,59,62–64^.

For example, consider the experience of a girl spending a day at the lake with her father (Fig. 5j, k). It may contain predictable relationships about birds flying, swimming, and perhaps even catching fish, as well as predictable relationships about fresh-picked strawberries tasting sweet. Our theory posits that these relationships should be extracted from the experience and integrated with memories of related experiences, through regulated systems consolidation, to produce, reinforce, and revise predictions (generalizations). On the other hand, unpredictable cooccurrences, such as the color of her father’s shirt matching the color of the strawberries, should not be consolidated in the neocortex. They could nevertheless remain part of an episodic memory of the day, which would permanently depend on the hippocampus for retrieval.

Go-CLS highlights the normative benefits of complementary learning systems, reconciles previous experimental results (Supplementary Material 11), and makes testable predictions that could support or refute the theory (Supplementary Material 12). A critical insight from Go-CLS theory is that gradual consolidation of past experiences benefits generalization performance most when experience is limited and relationships are partially predictable (Fig. 4), mirroring ethologically realistic regimes experienced by animals living in an uncertain world^65^. This benefit occurs in a regime where the danger of overfitting is the highest^37,38,57,62–64^, highlighting the need for a regulated systems consolidation process.

Previous theories have also sought to reconcile these and other experimental observations. For example, multiple trace theory^20^ and trace transformation theory^21,22^ posit that episodic memories are consolidated as multiple memory traces, with the most detailed components permanently residing in the hippocampus. Contextual binding theory^19^ posits that items and their context remain permanently bound together in the hippocampus. These theories emphasize the role of the hippocampus in permanent storage of episodic details^19–21,66,67^, with the neocortex storing less detailed semantic components of memories. In contrast, Go-CLS posits that predictability, rather than detail, determines consolidation. Similarly, Go-CLS favors predictability over frequency, feature overlap, or salience as the central determinant of systems consolidation^68–70^.

The fact that an experience’s predictability is *a priori* unknown has important conceptual implications for regulated systems consolidation. Here we have shown that it’s sometimes possible to accurately infer predictability from data (Fig. 2). This capability allows accurate generalization that is likely critical for building high-fidelity models of the world. However, many studies suggest that the brain relies on suboptimal heuristics for decision making and other cognitive tasks, and regulating systems consolidation based on inaccurate heuristics could lead to mis-generalization and departures from the predictions of Go-CLS theory. For example, an interesting prediction of Go-CLS theory is that frequent misinformation should be consolidated less than rare gems from a wise source, but this prediction would fail if brains used frequency as a simple heuristic for predictability. Extreme mis-regulation of consolidation could relate to disorders such as PTSD^71^. Modeling regulated systems consolidation in real world scenarios thus requires a precise understanding of the brain’s predictability estimation algorithm. Targeted experimental tests of Go-CLS theory could avoid this issue by focusing on tasks where animals generalize accurately.

Go-CLS theory does not specify the biological mechanisms by which memory consolidation should be regulated. Given the prominent role of sequence replay in existing mechanistic hypotheses about systems consolidation, this would be a natural target for regulation^72–81^. One possibility would be that memory elements reflecting predictable relationships could be replayed together, while unrelated elements are left out or replayed separately. Another would be that entire experiences are replayed, while other processes (e.g., attention mechanisms enabled by the prefrontal cortex^82,83^) regulate how replayed events are incorporated into neocortical circuits that store generalizations^66^.

The proposed principle that the degree of predictability regulates systems consolidation reveals complexities about the traditional distinctions between empirically defined episodic and semantic memories^5^. Most episodic memories contain both predictable and unpredictable elements. Unpredictable coincidences in place, time, and content are fundamentally caused by the complexity of the world, which animals cannot fully discern or model. Memorizing such unpredictable events in the hippocampus is consistent with previous proposals suggesting that the hippocampus is essential for incidental conjunctive learning^54^, associating discontiguous items^84^, storing flexible associations of arbitrary elements^10^, relational or configural information^85^, and high-resolution binding^86^. However, our theory holds that predictable components of these episodic memories would consolidate separately to form semantic memories that inform generalization. We anticipate that psychologists and neurobiologists will be motivated by the Go-CLS theory to test and challenge it, with the long-range goal of providing new conceptual insight into the organizational principles and biological implementation of memory.

## ACKNOWLEDGMENTS

The authors thank Tim Behrens, Brad Hulse, David Kastner, Jay McClelland, Brett Mensh, Sandro Romani, and three referees for helpful comments on the manuscript. We thank Julia Kuhl for illustrations and helpful discussions of the figures. This work was supported by the Howard Hughes Medical Institute (WS, NS, JEF) and the Janelia Visiting Scientist Program (AS). MA and AS were additionally supported by the Swartz Foundation, and AS by a Sir Henry Dale Fellowship jointly funded by the Wellcome Trust and the Royal Society (Grant Number 216386/Z/19/Z), the Gatsby Charitable Foundation, and the CIFAR Azrieli Global Scholars program.

## METHODS

### Teacher-Student-Notebook framework

Please refer to the Supplementary Material for a detailed description of the Teacher-Student-Notebook framework. The following sections provide a brief description of the framework and simulation details.

#### Architecture

The teacher network is usually a linear shallow neural network generating inputoutput pairs {*x^μ^, y^μ^*}, *μ* = *P*, through 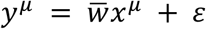, as training examples. Components of the teacher’s weight vector, 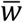, are drawn *i.i.d.* from 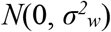; components of the teacher’s input patterns, *x^μ^,* are drawn *i.i.d.* from *N*(0, 1/*N*), where *N* is the input dimension; *ε* is a Gaussian additive noise drawn *i.i.d.* from 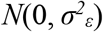. The signal-to-noise ratio (SNR) of the teacher’s mapping is defined as 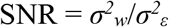, and we set 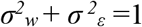 to generate output examples of unit variance. For the simulations in Figs. 2, 3, and 4, the student is a linear shallow neural network whose architecture matches the teacher (both with input dimension = 100 and output dimension = 1). We relaxed this requirement in Fig. 5 to allow mismatch between the teacher and student architectures (see **Generative models** section below). Components of the student’s weight vector, *w*, are initialized as zeros *(i.e., tabula rasa),* unless otherwise noted. The notebook is a sparse Hopfield network containing *M* binary units (states can be 0 or 1, *M* = 2000 to 5000 unless otherwise noted). The input and output layers of the student network are bidirectionally connected to the notebook with all-to-all connections.

#### Training procedure

Training starts with the teacher network generating *P* input-output pairs, with certain predictability (SNR), as described above. For each of these *P* examples, the teacher activates the student’s input and output layers via the identity mapping; at the same time, the notebook randomly generates a binary activity pattern, *ξ^μ^, μ* = 1,⋯, *P*, with sparsity *a*, such that exactly *aM* units are in the “1” state for each memory. At each of the example presentations, all of the notebook-to-notebook recurrent weights and the student-to-notebook and notebook-to-student interconnection weights undergo Hebbian learning (Supplementary Material 1). This Hebbian learning essentially encodes *ξ^μ^* as an attractor state and associates it with the student’s activation {*x^μ^, y^μ^*}, for *μ* = 1,⋯, *P*.

After all *P* examples are encoded through this one-shot Hebbian learning, at each of the following training epochs, 100 notebook-encoded attractors are randomly retrieved by initializing the notebook with random patterns and letting the network settle into an attractor state through its recurrent dynamics. Notebook activations are updated synchronously for 9 recurrent activation cycles, and we found that each memory was activated with near uniform probability. Once an attractor is retrieved, it activates the student’s input and output layers through notebook-to-student weights. Since the number of patterns is far smaller than the number of notebook units (*P* << *M*) in our simulations, the Hopfield network is well below capacity, and most of the retrieved attractors were perfect recalls of the original encoded indices. However, real hippocampal networks exhibit active forgetting that may enhance generalization or memory capacity^23,87^, and it would be interesting to consider alternate notebook models that incorporate forgetting effects^88^. Reactivation of the student’s output through the notebook, 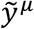, is then compared to the original output activated by the teacher, *y^μ^*, to calculate how well the reactivation resembles the original experience, quantified as the mean squared error. For error corrective learning, the student uses the notebook reactivated 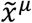 and 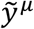. By comparing the student output that is generated by the reactivated 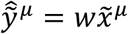 and the reactivated student output 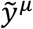 for all *P* examples, the student updates *w* using gradient descent with 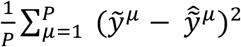 as the loss function. The weight update follows:

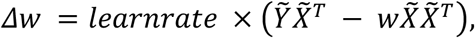

where 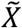 and 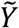 are the column wise stacked matrix form of the 100 reactivated input and output data points, respectively. Training continues for 500-5000 epochs, and *learnrate* ranges from 0.005 to 0.1. In our simulations, as long as the *learnrate* is sufficiently small (0.1 or smaller), the results stay qualitatively constant, and the main results do not depend on the specific choices of *learnrate. P_test_* number of additional teacher-generated examples, typically 1000, are used to numerically estimate the generalization error at each time step by 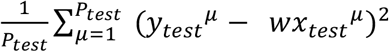. For some simulations we have applied optimal early-stopping regularization, where we stop the training when the generalization error reaches minimum.

#### Retrograde amnesia curves

We draw the following connections from network performance in terms of mean squared error to memory and generalization scores, which is typically measured by behavior responses in a task designed to test memorization or generalization performances. At the time when no training occurs, the network error corresponds to a random performance in a task, which is typically set as the zero of a memory retrieval metric. As the error decreases with training, the error is related to the memory retrieval score as follows: score = (*E_0_* - *E_t_*)/*E_0_*, where *E* stands for memorization error or generalization error, and the subscript indicates the timestep during training. This is stating that the memory retrieval score at each time point is negatively correlated to the error at that time and normalized into a range where 0 indicates chance performance and 1 indicates perfect performance. During memory retrieval (or generalization), the system chooses whichever available module with lower memorization error (or generalization error). To simulate notebook lesioning at time *t*, the system starts to use only the student for memory recall, in addition, student’s memory score will remain unchanged with time due to the lack of notebook-mediated systems consolidation. In Fig. 3e, both the SNR and amount of prior learning were varied to produce the diverse shapes of retrograde amnesia curves. For the control simulation, SNR is set to ∞. For the solid lines of retrograde amnesia curves, SNR values are 0.01, 0.1, 0.3, 1, and 8. SNR is set to 50 for the dotted lines simulating the effect of prior consolidation. Each line is a different simulation with the amount of prior consolidation ranging from 8 epochs to 2000 epochs (*learnrate* = 0.005). *N* = 100 and notebook size *M* = 5000. *P* = 100 for the varying SNR simulations and *P* = 300 for varying prior consolidation simulations.

#### Generative models for diverse teachers

To explore different ways unpredictability can exist in the environment, we generalize the teacher-student-notebook framework by relaxing the linear and size matched settings to allow for more complex teachers as generative models for producing training data. For the nonlinear teacher setting, a nonlinear activation function is applied to the linear transformation generate the teacher’s output. A sine function was chosen for the simulation in Fig. 5e. The corresponding noisy teacher’s SNR is numerically determined from the complex teacher’s nonlinearity detailed in Supplementary Material, Section 10. For the partially observable teacher, the input layer is larger than the student’s, and the student can only perceive a fixed subregion of the teacher input layer. The exact size of the partially observable teacher is set to match the calculated equivalent SNR of the complex teacher.

#### Data availability

No data were collected for this theoretical study.

#### Code availability

Code reproducing the results is available at https://github.com/neuroai/Go-CLS_v1

## Supplementary Material

### 1 The Teacher-Student-Notebook framework

We consider a setting in which an agent experiences examples of a relationship in the environment in the form of *P* pairs of activity patterns {*x^μ^, y^μ^*}, *μ* = 1, ⋯, *P*. Given the *input* activity vector of dimension *N, x^μ^* ∈ *R^N^*, the agent must both memorize the associated scalar *output* activity *y^μ^* and develop the ability to predict outputs for new inputs. For example, *x^μ^* might represent activity in visual cortex in response to an event like seeing a bird, and *y^μ^* might represent activity in a higher association cortex derived from a caregiver’s speech: “Look, a bird!” The agent wishes to memorize what was seen and said in this specific instance, as well as learn what birds look like more generally.

For any given event, there will be many such relationships to learn, which collectively encode many diverse features in the environment. For instance, while viewing the bird, other neural circuits may encode the spatial location of the event, the time of day, other objects in the scene, and so on. Our model first considers just one of these relationships, and we return to having multiple relationships in the later sections of this document. Our theory contains three neural network components, and we refer to it eponymously as the teacherstudent-notebook framework. We now describe each of these components in sequence.

#### Teacher Network

The ground truth relationship between inputs and output is represented by a *teacher* network. It generates input-output pairs by first drawing an input vector, x, in which each element is *i.i.d.* normal with variance *1/N*, i.e. 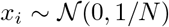, *i* = 1, ⋯, *N*. Thus, the norm of the input vector is one in expectation. Next, the teacher labels this input according to the rule

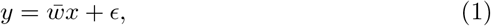

where 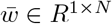 are the teacher weights, and *ϵ* is the teacher output noise. In words, the teacher is simply a shallow linear network with output noise. We take the teacher weights to be *i.i.d.* normal with variance 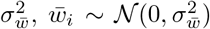, *i* = 1, ⋯ , *N*, and the output noise is *i.i.d.* normal with variance 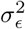. Note that the noise varies across examples, but the weights are fixed for all examples.

A key parameter of this setting is the *signal-to-noise* ratio (SNR),

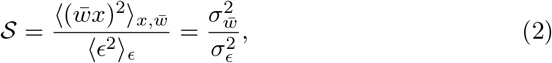

where 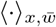 denotes the average over input patterns and teacher weights, and 〈·〉 denotes the average over noise. This ratio measures the extent to which the teacher’s output follows a systematic mapping between input and output. To fix the scale across different teachers, we consider the case where the variance of the teacher’s output is one,

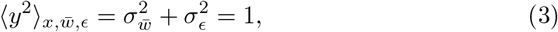

such that the SNR fixes the variances,

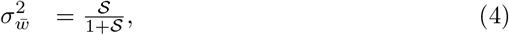

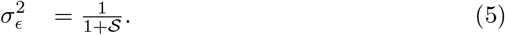

Conceptually, the teacher provides a generative model of the environment. We emphasize that taking the teacher to be a simple neural network does not reflect an assumption that the real environment is either simple or a neural network. Rather, the teacher network can be thought of as containing the optimal synaptic weights for approximating the true generative model of the environment, which may reflect diverse causal processes, from the physics of the world to the neural circuits that generate input and output activity patterns. In this sense, the teacher is a useful abstraction, not a mechanistic theory of the environment. We discuss further interpretations of the teacher in Section 10.

#### Student Network

The goal of the *student* network is to learn to approximate the relationship defined by the teacher. Here we take the student to have the same architecture as the teacher, that is, it is a shallow linear network that receives an *N*-dimensional input, *x*, and produces a predicted output, 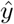, according to

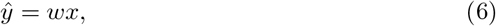

where *w* ∈ *R*^1×*N*^ are the student weights. These weights are learned using gradient descent on a loss function 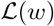

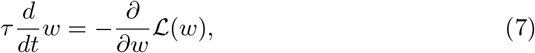

here formulated in continuous time (also known as gradient flow) with time constant *τ*. We take the loss function to be the mean squared error over the example patterns,

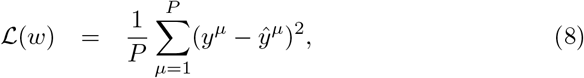

where *μ* = 1, ⋯ , *P* indexes examples, *y^μ^* is the scalar target output, and 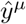 is the scalar output prediction in response to input vector *x^μ^*. As described in more detail subsequently, the target patterns that drive learning can have multiple sources-they may come directly from the teacher or from a memory of past examples.

The performance of the student can be measured in two ways. First, its predictions can be evaluated on the specific examples *μ* = 1, ⋯ , *P* seen during training, which we refer to as the *memory* error 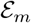 (also known as the *training error* in machine learning contexts),

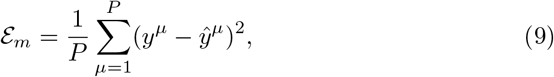

where here we have not indicated the time dependence of 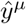 for notational simplicity. Second, the student’s predictions can be evaluated on novel inputoutput pairs drawn from the teacher, which we refer to as the *generalization* error *E_g_* (also known as the *test error* in machine learning contexts),

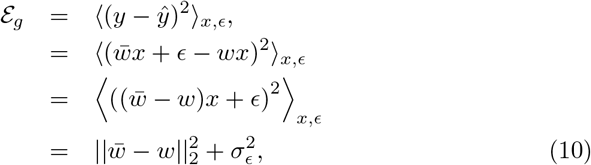

where 〈·〉_*x,ϵ*_ denotes the average over the teacher input distribution and output noise distribution, and we have used the fact that these distributions are independent.

#### Notebook Network

Finally, the job of the *notebook* network is to faithfully memorize experienced patterns as attractors of neural network dynamics, making possible later recall and replay. We consider a notebook of *M* neurons recurrently connected through the *M* × *M* weight matrix *J*. The activity in this network is binary, *h* ∈ {0,1}^*M*^, and evolves according to

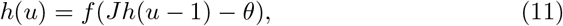

where *f* is the Heaviside step function, *u* denotes discrete time steps of synchronous activity propagation, and *θ* is a threshold that can have a fixed value or be dynamically adapted to maintain a desired sparsity of activity (as described subsequently).

The notebook represents memorized patterns as binary vectors of zeros and ones by embedding these vectors as fixed points of the dynamics in Eq. (11). In particular, to store input-output pairs, {*x^μ^, y^μ^*}, *μ* =1, ⋯ , *P*, the notebook first chooses *P* binary (0/1) vectors of length *M*, uniformly at random from the set of vectors with sparsity *a* (i.e. with exactly *aM* nonzero entries). These binary patterns of activity in the notebook act as distinctive neural codes to be associated with each pattern, and this notebook code is sometimes referred to as a memory index.

Stacking the binary patterns into the columns of the *M* x *P* matrix *ξ*, and similarly stacking the input and output patterns into the *N* x *P* and 1 x *P* matrices *X* and *Y* respectively, the weights within the notebook and between the notebook and student are given through a Hebbian scheme,

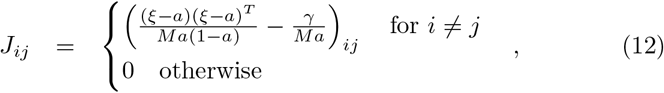

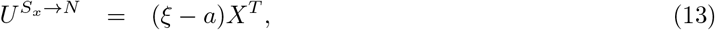

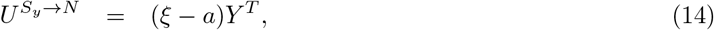

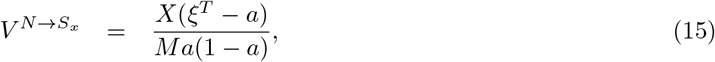

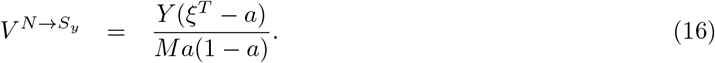

Here *U^S_x_→N^* ∈ *R^M×N^* and *U^S_y_→N^* ∈ *R*^*M*×1^ map from the student inputs *x* and output *y* to the notebook activity *h*, and the matrices *V^N→S_x_^* ∈ *R^N×M^* and *V^N^→^S_y_^* ∈ *R*^1×*M*^ perform the reverse mapping from the notebook activity back to the student input and output. For simplicity and tractability, we take all student neurons to be linear. The parameter *γ* in Eqn. 12 implements global all-to-all inhibition, which causes activity that is far from stored patterns to decay to a silent state [11]. In simulations, we take *γ* = 0.6, which lies in the range theoretically derived to stably store the intended patterns without spurious attractors in this model [11]. These pathways allow diverse interactions between notebook and student, and we describe a number of specific interaction patterns subsequently.

The mean subtraction and normalization in these updates have been chosen to aid performance, as derived subsequently in Section 5.1 for connections from notebook neurons. In essence, the notebook generates distinct, pattern-separated activity patterns, stabilizes these as attractors of its recurrent dynamics, and links these bidirectionally to the student’s input and output neurons to facilitate later replay and reactivation.

### 2 Learning setting

The teacher-student-notebook framework can allow for diverse learning settings in which examples from the teacher arrive at different times and in different quantities. Here we usually characterize memorization and generalization performance in a simple setting: the *single-batch, high-dimensional* regime. That is, we consider a scenario where an organism receives *P* training experiences up front in a short time window, and memory and generalization performance are evaluated subsequently over longer periods of time. For instance, a human subject might learn a task in a single hour long session but then be tested after several weeks’ delay, or a rodent might perform several trials in a water maze on one day and be tested on the next. In our framework, these *P* experiences are drawn *i.i.d.* from the teacher and constitute one single batch for learning and consolidation. For convenience, we can collect this batch of samples into the *N* x *P* matrix *X* with columns *x^μ^, μ* = 1, ⋯ , *P*, and the 1 x *P* row vector *Y* with elements *y^μ^, μ* =1, ⋯ , *P*.

Given abundant training experience (*P* >> *N*), many different learning schemes can converge to similar performance. However, real world learning is often severely data limited. Animals may receive only one or two foot shocks. A human subject may need to learn a new visual discrimination (possibly dependent on millions of pixels) from just a few blocks of training trials. Real world settings therefore place a premium on learning from limited experience. Moreover, neuronal networks in the brain are typically very large relative to the amount of training experience. Even a simple visual discrimination may engage a network of millions or billions of neurons interconnected by billions or trillions of adjustable synapses. To address this large network, limited data setting, we analyze the *high-dimensional* regime, in which the size of the student network and the number of training samples both tend to infinity (*N* ! ∞, *P* ! ∞), but their ratio *α* = *P/N* remains finite. The *load parameter α* is a key parameter of our setting, and it measures the amount of experience relative to the number of tunable synapses in the student network. For *α* < 1, the network has more tunable parameters than training experiences, allowing analysis of highly overparametrized learning settings. For *a* >> 1, the network has many more training experience than tunable parameters, reflecting the more standard classical regime of statistics.

While in this paper we emphasize this single-batch, high-dimensional learning setting, future work in the teacher-student-notebook framework could investigate more complex scenarios where examples continue to arrive over time.

### 3 Interaction policies & performance

The single-batch learning setting still allows diverse possible interaction policies between the modules in the teacher-student-notebook framework. These interaction policies specify which modules undergo learning, from what activity patterns (e.g. from the teacher, or from replay from the student), and which modules are used to answer queries for new experiences. We consider four interaction policies, meant to typify common approaches to learning and consolidation.

**Online Student.** Only the student is trained, without any replay. Each example drives one update of error-corrective learning and is never revisited. This strategy provides a reference point for performance of a system based on online gradient descent learning.
**Online Notebook.** Only the notebook is used. Each example is stored in the notebook with Hebbian updates, and predictions for novel inputs are generated using the notebook only. This strategy provides a reference point for performance of a system based on Hebbian memorization, without replay-guided learning.
**Memory-optimized Replay.** This strategy initially stores all experiences in the notebook and trains the student using notebook-driven reactivations until the student has fully memorized all examples. This is similar to the standard theory of systems consolidation.
**Generalization-optimized Replay.** This novel strategy, proposed in this work, initially stores all experiences in the notebook but only trains the student using notebook-driven reactivations as long as generalization performance improves.

The next four sections of the supplement sequentially characterize the memorization and generalization performance of each of these interaction policies.

### 4 Online Student Policy

In the online student policy, each example *x^μ^* ∈ *R^N^, μ* = 1, ⋯ , *P*, in the batch is visited in order and a single step of error corrective gradient descent learning is applied with an example-dependent learning rate *η^μ^*. In this section we characterize the expected generalization error dynamics under this scheme; to ensure a robust normative comparison to other policies, we derive the globally optimal learning rate function that maximizes generalization performance after all updates.

#### 4.1 Generalization dynamics with example-dependent learning rate

Upon receiving each example *μ* = 1, ⋯ , *P*, the student weights are updated according to

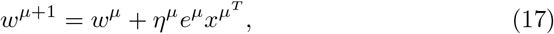

where *w*^*μ*+1^ is the weight vector resulting from the *μ*th learning step, *η^μ^* is the learning rate of this step, *x^μ^* is the *μ*th input example, and 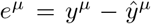 is the error between the network’s output and the target output for this example. We assume that the initial weights are zero, *w*^1^ = 0. Using the teacher model, 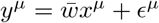, we have

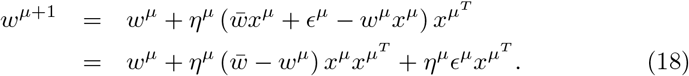

In contrast to Eqn. (10), which expresses the generalization error 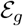 for a specific student and teacher, here we ask what the expected generalization error is for a randomly drawn teacher by averaging over the teacher weight distribution as well. That is, we track the expected generalization error 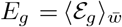, where the average is over the teacher weight distribution. In the highdimensional regime, the generalization error is self-averaging, such that any specific realization closely tracks this expected generalization error, as will be verified by a close match between single simulations and the average dynamics we derive. The expected generalization error before example *μ* is

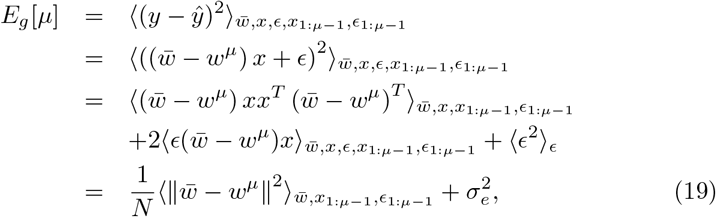

*x*_1:*μ*_ and *μ*_1:*μ*_ denote history of training patterns and corresponding additive noise, respectively. We used that 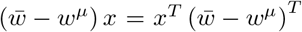 is a scalar, that *ϵ* is zero mean and independent of all other terms, and that *x* is multivariate normal with covariance matrix 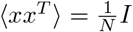. Similarly, note that after example *μ*, the expected generalization error becomes

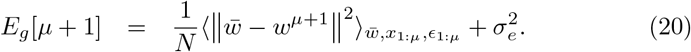

Substituting in Eqn. 18, we have

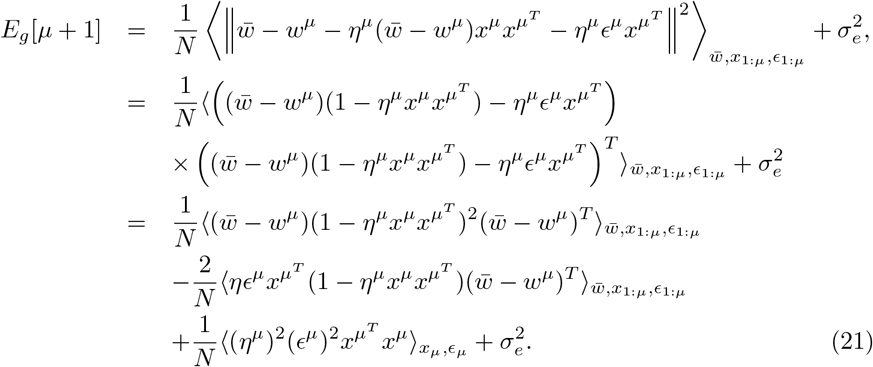

The term linear in the noise again vanishes, and we note that 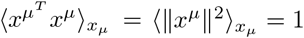. Therefore, the last term’s expectation is 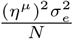, and

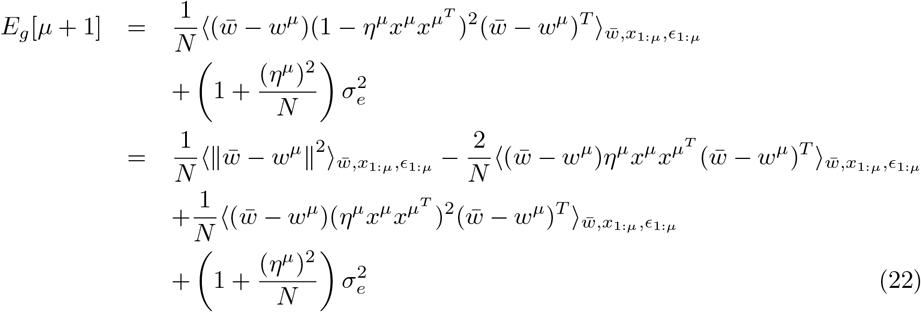

To simplify this expression, it’s convenient to note that

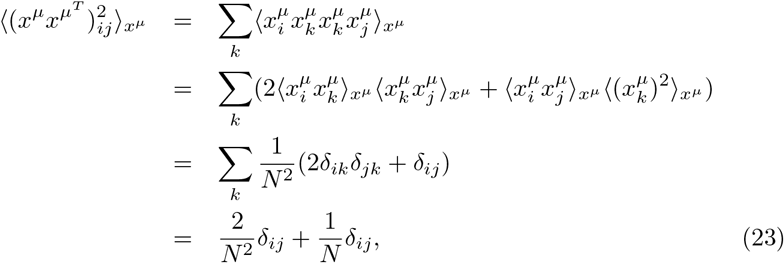

where the second line follows from Wick’s theorem for Gaussian moments, and *δ_ij_* is the Kronecker delta. This implies 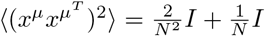, and together with 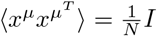, our expression for *E_g_*[*μ* + 1] becomes,

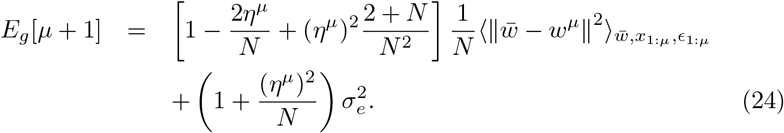

This weight norm is related to the generalization error by Eqn. 19, which enables a recursive equation for the generalization error

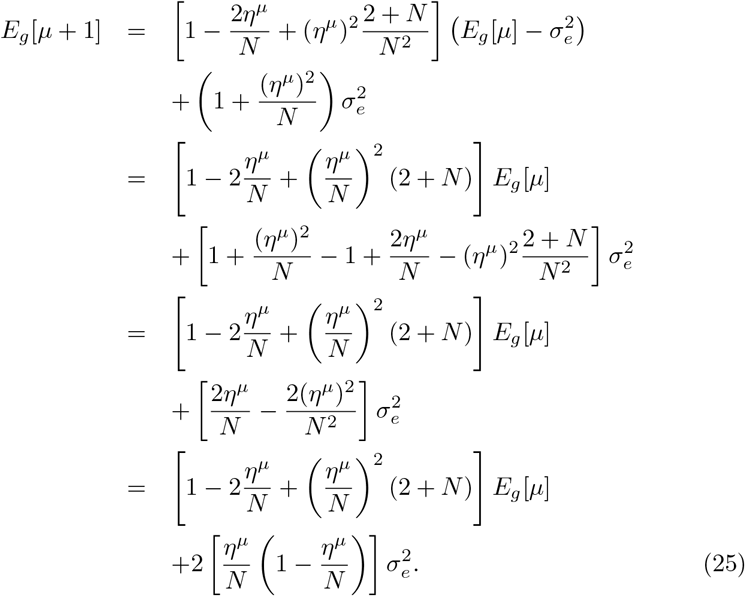

Now passing to the limit *N* >> 1, we have

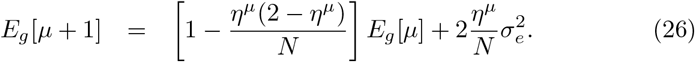

We then enter the high-dimensional regime where *α* = *μ/N* and consider the new continuous variables *E_g_*(*α*) ≈ *E_g_*[*αN*] for the generalization error and *η*(*α*) ≈ *η^αN^* for the learning rate^1^. We wish to calculate an equivalent differential equation,

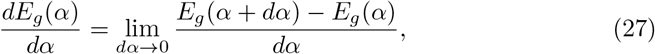

where we take *dα* = 1/*N*, which is infinitesimal in the limit *N* → ∞, to approximate the increment provided by a single new example. Thus

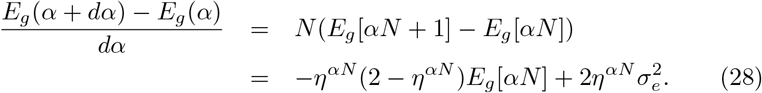

We thus have the ordinary linear differential equation

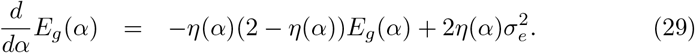

The solution can be found through the method of integrating factors. In particular, we define

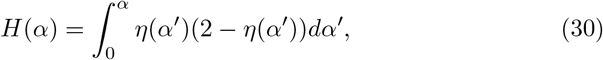

and find

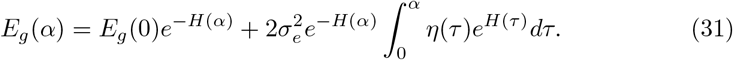

#### 4.2 Optimal online learning rate

Equation (31) yields the expected generalization error for arbitrary learning rate functions. To ensure a fair normative comparison to other methods, we now compute the optimal learning rate as a function of example. We again begin by considering a discrete sequence of examples, and we will take the highdimensional limit at the end.

Let *η^*,μ^* denote the learning rate schedule that minimizes the expected generalization error on example *T* = *αN*. Also let

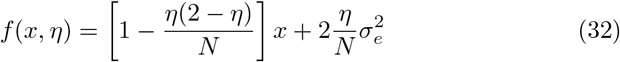

be the discrete dynamics update from Eqn. (26), that is, the generalization error on example *μ* + 1 if the generalization error on example *μ* is *x* and the learning rate used on example *μ* is *η*.

At the penultimate example before the deadline, *T* – 1, because there is only one update left, the best learning rate is given by greedily optimizing *f*,

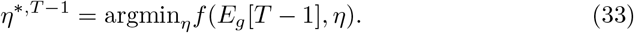

We directly perform the minimization by differentiating with respect to *η* and setting this derivative to zero,

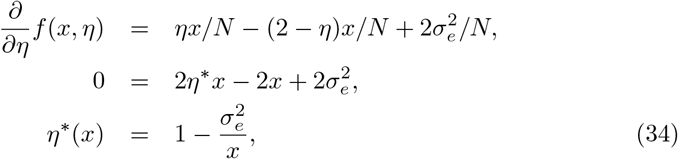

which yields the optimal update of

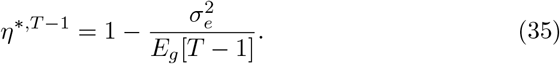

The final generalization error as a function of the penultimate generalization error, *x*, is thus

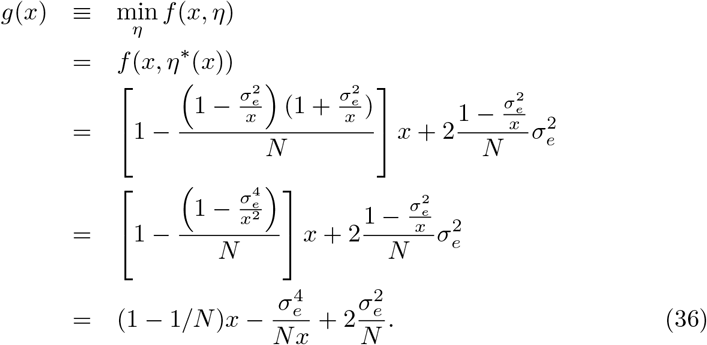

Differentiating with respect to *x*, we have

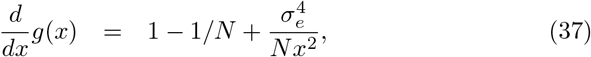

which is strictly positive for *N* ≥ 1, *x* > 0. This indicates that the function *g*(*x*) is strictly increasing, meaning that larger generalization errors at the penultimate step directly translate into larger generalization errors at the deadline.

Let *v_μ_*(*x*) denote the optimal final generalization error on example *T*, starting from an error of *x* at step *μ* and choosing the optimal learning rate thereafter. We have shown that *v*_*T*–1_(*x*) = *g*(*x*), and it is strictly increasing. Now for the inductive step, assume that *v*_*μ*+1_(*x*) is strictly increasing. Then

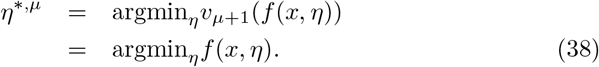

Therefore the optimal learning rate is again selected by greedily minimizing *f*(*x, n*). Finally, we note that *v_μ_*(*x*) = *v*_*μ*+1_(*g*(*x*)) is the composition of strictly increasing functions, and therefore strictly increasing. This establishes the inductive hypothesis and yields the optimal learning rate function for all examples

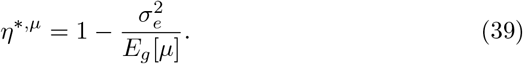

In the high-dimensional regime, the optimal learning rate is thus

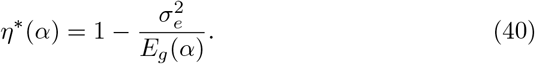

Inserting this optimal learning rate function back into Eqn. (29) yields the following optimal generalization error dynamics,

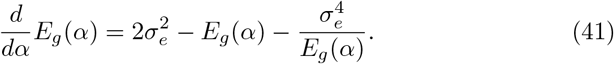

### 5 Online Notebook Policy

In the online notebook policy, each example is stored in the notebook according to the Hebbian scheme in Eqns. (12)-(16). The notebook is then used to make predictions even for novel inputs, by allowing the notebook to converge to an attractor and reading off the predicted output.

In particular, an input *x* arriving at the student from the teacher can be used to seed recurrent pattern completion in the notebook, by letting *h*(0) = *f*(*U*^*S_x_*→*N_x_*^) and then running the notebook dynamics. In the simulations in the main text, rather than run the recurrent dynamics to convergence, we use the pattern obtained after 9 updates. At each update, the neurons are ranked by net input and the threshold *θ* is chosen so that the top *aM* are active (in the case of ties, slightly more neurons can be active). After the network dynamics have settled on some pattern 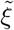, a predicted output can be generated (using just the notebook) as 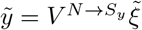.

This section shows that, in the high-dimensional setting considered here, the notebook attains low memorization error (i.e. error on already-experienced examples) but is incapable of generalization.

#### 5.1 Hebbian learning rule scale factor and offset

The memorization ability of recurrent attractor networks, as well as the performance of Hebbian plasticity rules in mapping from notebook activity patterns to student activity patterns, is known to depend on the statistics of the patterns and the specific form of the learning rule used to configure the weights [18, 5, 6, 33]. We begin by justifying the scaling and subtractive offsets in Eqns. (12)-(16), typically as an approximate implementation of the pseudoinverse learning rule given our sparse pattern statistics.

##### 5.1.1 Recurrent weights

The job of the notebook is to faithfully memorize example patterns as attractors of neural network dynamics. The pseudoinverse learning rule is a flexible mechanism to memorize these patterns, wherein the *M* × *M* matrix of recurrent notebook connections would be

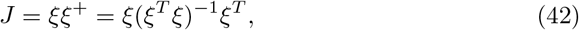

where *ξ*^+^ is the pseudoinverse of *ξ*, and we assumed that *P* ≤ *M*. Suppose that the neural network dynamics have the form *h*(*u*) = *f*(*Jh*(*u* – 1)), where *h* is the pattern of notebook activity. Assuming that *f*(0) = 0 and *f*(1) = 1 (e.g. *f* may be linear, threshold-linear, or binary), then these weights would successfully memorize all *P* patterns as steady-states of the network dynamics. In particular, note that

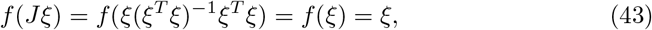

so that the network dynamics map each memorized pattern back onto itself^2^. It is instructive to expand the pseudoinverse weights in terms of the stored patterns,

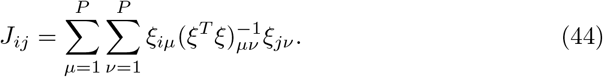

This reveals a practical problem with the pseudoinverse learning rule, as the storage prescription for each pattern depends on the other stored patterns through the inverse pattern correlation, 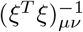.

The Hopfield model can be viewed as a solution to this problem that assumes simple random statistics for *ξ* in order to simplify the necessary structure of the learning rule. In particular, suppose that each memory randomly assigns aM neurons to the 1-state and (1 – *a*)*M* neurons to the 0-state. Thus, a quantifies the fraction of 1-states in the memorized patterns, and we refer to a as the sparseness parameter. We also assume that the memorized patterns are statistically independent from each other. These statistics imply that

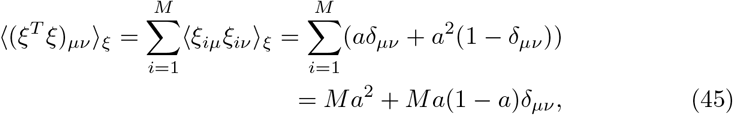

where 〈·〉 now denotes the average over notebook patterns. In matrix notation, this implies that

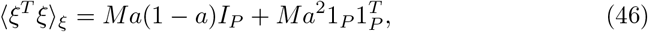

where *I_P_* is the *P* × *P* identity matrix, and 1_*P*_ is the *P*-vector of ones. This form allows us to use the Sherwood-Morrison formula,

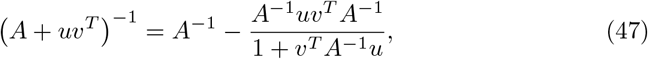

with *A* = *Ma*(1 – *a*)*I_P_, u* = *Ma*^2^1_*P*_, and *v* = 1_*P*_ to obtain

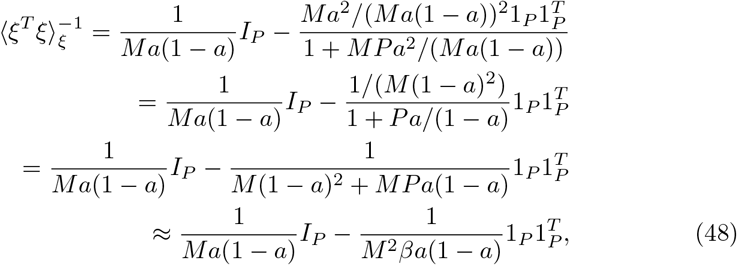

where the final approximation used *P* = *βM*, *β* = *O*(1), and *M* ≫ 1. The Hopfield model approximates the pseudoinverse learning rule by replacing (*ξ^T^ξ*)^-1^ by 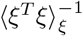. To see what this means, we need to do a bit more algebra:

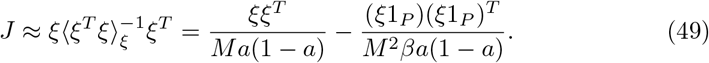

The Hopfield model also approximates *ξ*1_*P*_ by 〈*ξ*1_*P*_〉 = *Pa*1_*M*_, where 1_*M*_ is the *M*-vector of ones, such that

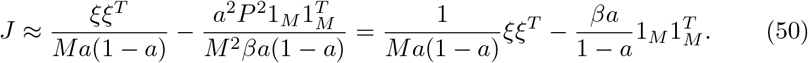

To compare this to the Hopfield model, we first consider a general Hebbian weight matrix of the form,

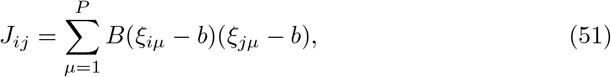

where *B* and *b* are constants that scale and center the learning rule. Again using the approximation that *ξ*1_*P*_ ≈ 〈*ξ*1_*P*_〉_*ξ*_ = *Pa*1_*M*_, we find

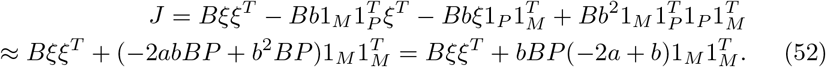

Comparing Eqs. (50) and (52), we see that the two correspond when

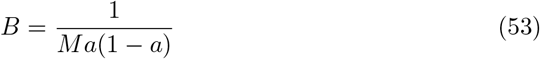

and

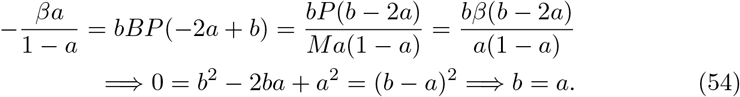

Therefore, the pseudoinverse rule can be approximated by the Hebbian rule,

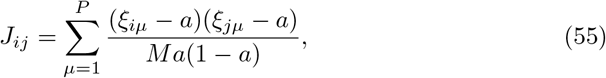

which is the weight matrix of the Hopfield model and the first term in Eqn. (12) of the notebook learning rules.

##### 5.1.2 Notebook-to-student weights

Similar to the Hopfield storage prescription used to store binary indices as fixed points of the recurrent notebook dynamics, here we assume Hebbian connectivity between the notebook and student. In particular, we can form the (*N*+1) ×*P* matrix *Z* by vertically stacking the matrices *X* and *Y*, such that *Z* represents the combined student input-output activity to be stored. We also define the (*N* +1) x *M* matrix *V* by vertically stacking the matrices *V*^*N*→*S_x_*^ and *V*^*N*→*S_y_*^, which represents the mapping from notebook activity to student activity. In this setting the relevant pseudoinverse learning rule for the weights from notebook to student neurons is

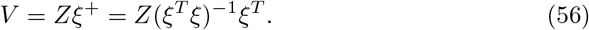

The same approximations used in the previous section lead to

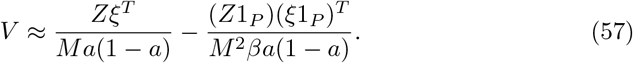

Replacing *ξ*1_*P*_ by 〈*ξ*1_*P*_〉 = *Pa*1_*M*_, we find

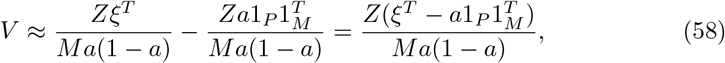

or

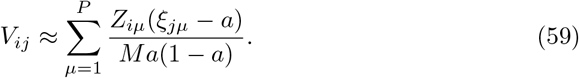

This is the Hebbian learning rule that we use to connect the notebook to the student for purposes of pattern reactivation (Eqns. (15)-(16)).

##### 5.1.3 Student-to-notebook weights

We did not derive the student-to-notebook weights from a pseudoinverse rule. In particular, the associated pseudoinverse rule would depend on *Z*^+^, which must be computed differently depending on whether *P* ≤ *N* +1 or *P*>*N* + 1. In contrast, we assumed that *P* ≤ *M* throughout, which allowed a unified expression for *ξ*^+^. Moreover, memory recall means that the notebook will sometimes be activated by a subset of student neurons, so defining weights based on *Z*^+^ may be inappropriate in some circumstances.

Nevertheless, the form of the student-to-notebook weights is justifiable. Define the *M* x (*N* + 1) matrix *U* by horizontally stacking the matrices *U*^*S_x_*→*N*^ and *U*^*S_y_*→*N*^, which represents the mapping from student activity to notebook activity. It’s useful to note that Eqs. (13)-(14) imply

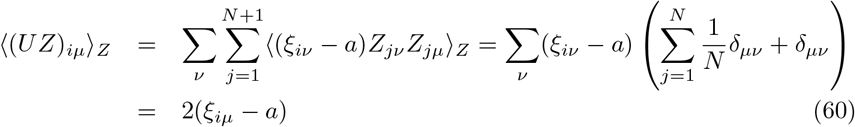

where the expectation is now over student patterns, and we noted that 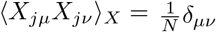 and 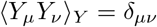. Therefore, 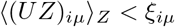, and 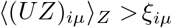 if *ξ_iμ_* = 1 and *a* < 0.5. Consequently, these weights are expected to seed the appropriate pattern in the binary notebook network with sparse memories. Similarly,

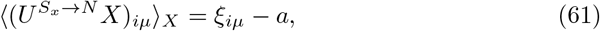

so 〈(*UX*)_*iμ*_〉*X* < *ξ_iμ_* if *ξ_iμ_* = 0, and 〈(*UX*)_*iμ*_〉*X* > *a* if *ξ_iμ_* = 1 and *a* < 0.5. The input neurons are thus also expected to seed the appropriate pattern if 0 < *θ* < *a*, or if the threshold is dynamically chosen to maintain the desired spareness level.

#### 5.2 Notebook memory error

With these Hebbian learning prescriptions in hand, we now characterize their performance. In this section, we consider the typical memory error by examining the statistics by which the notebook reactivates stored patterns of student activity. Previous studies of the Hopfield model [5, 6, 33] imply that large notebooks can accurately recall each random index if the number of stored patterns does not exceed the capacity of the network *P_c_* = *β_c_M*. Here we assume that M 1 and *P* < *P_c_*, such that erroneous index retrieval by the notebook is rare. Once a notebook memory index is accurately retrieved by the notebook’s dynamics, the notebook can generate a predicted output using the Hebbian weights from notebook to student output (*V^N→S_y_^*). The memory error of the notebook can thus be approximated as the typical error of this prediction.

As in the previous section, let *Z* be a (*N* +1) x *P* matrix that groups together all input and output neuron responses for all memorized patterns. Then the notebook reactivated student pattern is

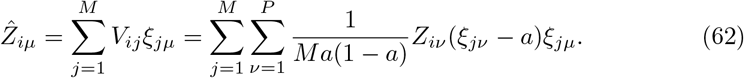

We first consider how well the notebook reactivates the student on average. In particular, averaging this expression over all possible notebook indices gives

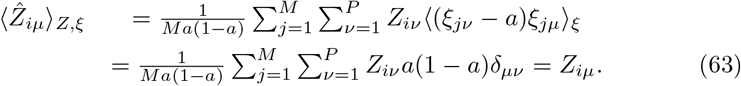

Therefore, the Hebbian learning rule is unbiased, and it on average reactivates all student neuron responses accurately.

However, the randomness of notebook indices does cause notebook-driven student reactivations to fluctuate away from these average values. To determine the magnitude of notebook memory error quantitatively, first note that the memory error of the notebook is

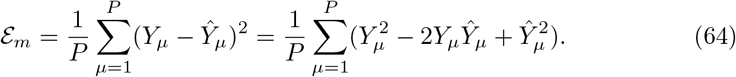

Averaging over possible notebook patterns, we find

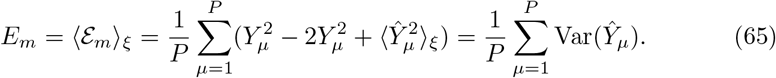

This variance term can be written

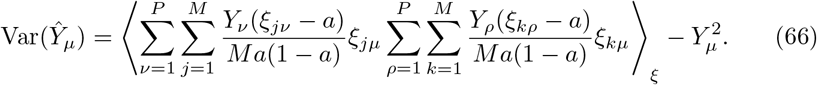

This expression shows that the exact value of the notebook training error depends on the specific realizations of the student outputs.

However, for practical purposes, it will be good enough to average Eq. (66) over possible student outputs, and noting that 〈*Y_μ_Y_v_*)_*Y*_ = *δ_μv_*, we find

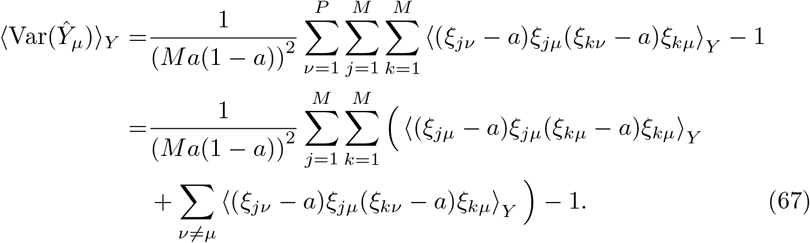

It is straightforward to evaluate the first expectation as

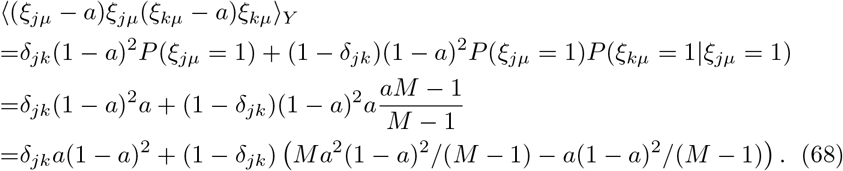

Because *μ* ≠ *v* in the second expectation of Eq. (67), it straightforwardly separates into the product of two terms:

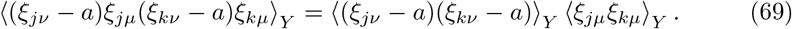

First,

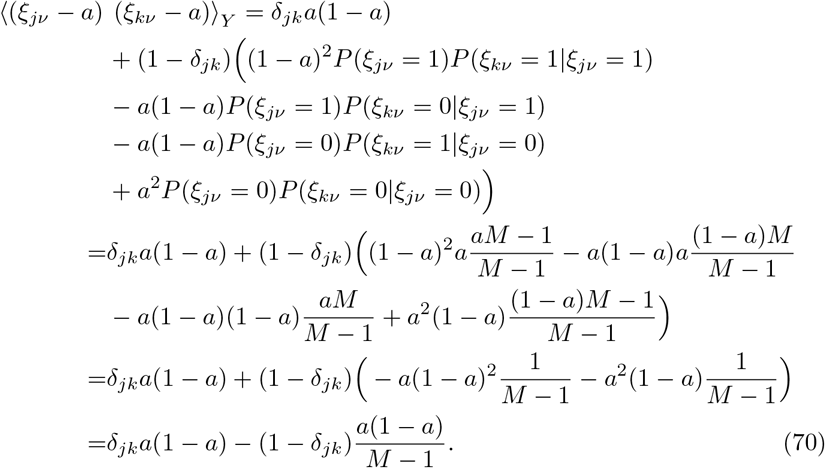

Second,

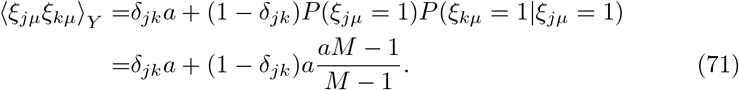

Combining these two terms, we find,

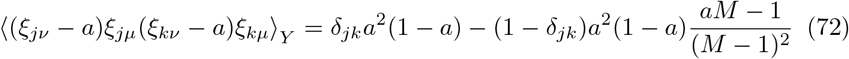

for *μ* ≠ *v*. Plugging these expressions back into the expression for 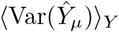, we find

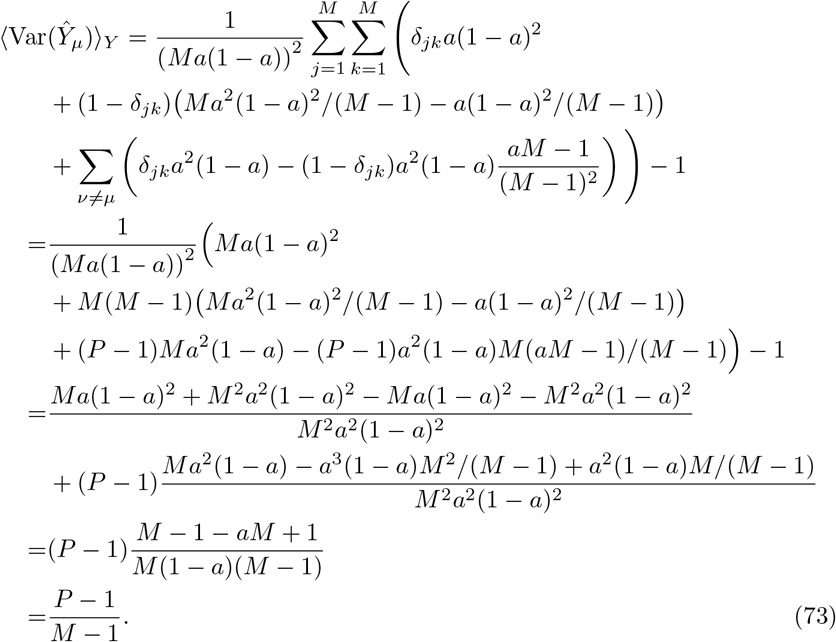

The proportionality of 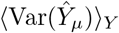 to *P* – 1 intuitively captures the interference of the Hebbian readout of memory *μ* from the other *P* – 1 memories that contribute to *V*^*N*→*S_y_*^.

Combining Eqs. (65) and (73), we find that the expected memory error of the notebook is simply

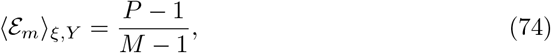

Remarkably, note that this expression is independent of the notebook’s sparseness. If *P* = *βM* and *M* ≫ 1, this implies that

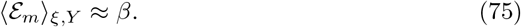

We thus see that the expected memorization error of the notebook scales with the number of memories stored in the system and can become significant when the loading is large. Note that this expression only makes sense if *β* < *β_c_*, because we’ve assumed faithful index reactivation within the notebook itself.

#### 5.3 Notebook generalization error

Next we examine the expected error when the notebook is used to predict the teacher output on a novel example. Because we operate the Hopfield network below capacity, it successfully embeds all patterns as fixed points with relatively large basins of attraction, and the previous section showed that the student reactivation error is modest. For simplicity, we therefore model the notebook as a nearest neighbor algorithm that operates by returning the output associated with the nearest stored pattern for any given input.

**Figure S1:**
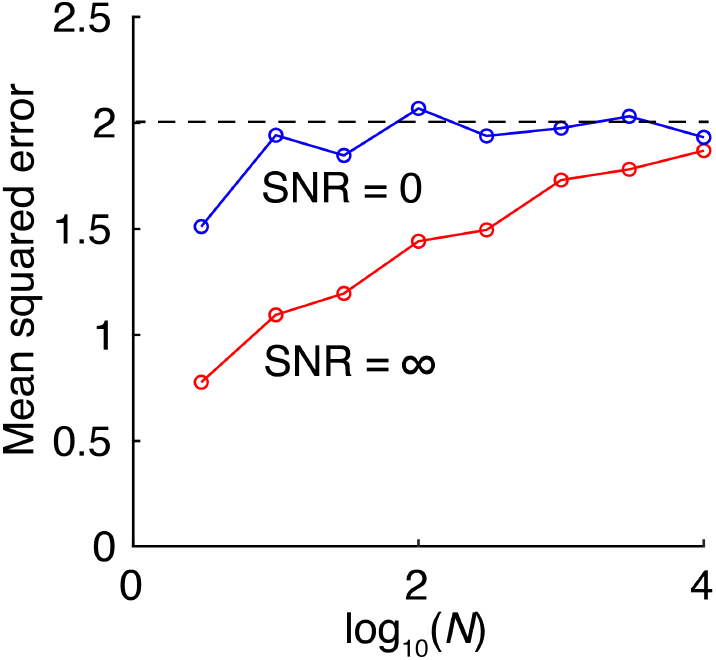
Numerical simulation of the generalization error of the nearest neighbor algorithm, for *α* = 1 and SNR 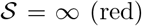 (red) and 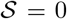 (blue). As input dimension N approaches infinity, generalization error in both cases approaches 2, consistent with our analytical derivation.

In particular, let *x^μ^, μ* = 1, ⋯ , *P* be the *N*-dimensional column vectors of stored inputs, and *y^μ^, μ* = 1, ⋯ , *P* be the associated outputs. For a novel input *x* 2 *R^N^*, we find the nearest neighbor as

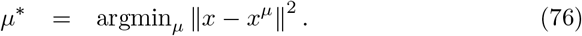

With the nearest neighbor identified, the prediction is 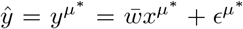. The expected generalization error is thus

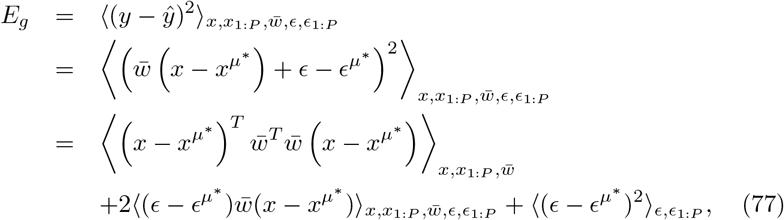

We used that 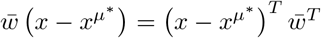 is a scalar. This form allows us to evaluate the expectation over teacher weights as 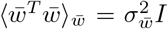. Also noting that the noise is uncorrelated with everything else, we find

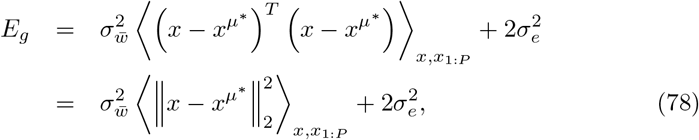

where the prefactor of 2 in the noise term reflects the independent fluctuations of *ϵ* and *ϵ^μ^*. We note that 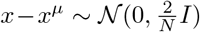 for all *μ*, and so 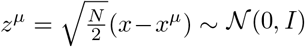. By the Gaussian Annulus Theorem (see e.g. Thm 2.9, pg 15 of [9]),

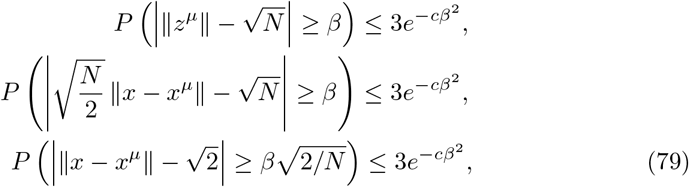

where *c* > 0 is a constant independent of *N*. By the union bound, the probability that one or more patterns among the *μ* = 1, ⋯ , *αN* fails to concentrate is no more than

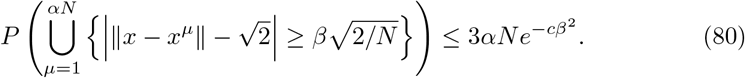

Therefore the minimum over all patterns will fail to concentrate with probability no more than

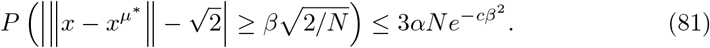

Choosing *β* = *N*^1/4^ we have,

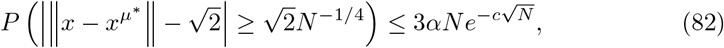

such that as *N* →∞, the minimum concentrates near 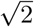 with probability one. Substituting back into the expression for the expected generalization error, in the high dimensional limit with high probability we have

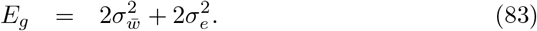

For our standard scaling where 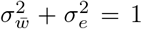, the error is therefore 2 regardless of the SNR. We note that this result applies in the high-dimensional limit where *N, P* → ∞ and their ratio is *α* = *P/N*. In finite size simulations, the generalization error can modestly differ, as shown in Fig. S1.

In essence, in the high-dimensional regime, the nearest neighbor is typically very far away from the new sample, such that generalization fails completely. In fact, it is so poor that always predicting zero would be better (attaining generalization error of 1 rather than 2 for our setting). This finding strongly motivates the need for a trained student, but we note that notebook-mediated generalization could be better in different settings where, for instance, input examples arise from a low number of clusters [15].

### 6 Memorization-optimized Replay Policy

In the memorization-optimized replay policy, each example is stored in the notebook according to the Hebbian scheme in Eqns. (12)-(16). These patterns can then be reactivated offline to drive learning. In the simulations reported in the main text, offline notebook reactivations undergo a two-step retrieval process:

1. A random binary pattern is used to seed the reactivation event. Starting at this random state, the notebook updates through the recurrent dynamics 9 times synchronously to retrieve a stored pattern. On each update, the threshold *θ* is chosen to enforce a sparsity of *a* (up to ties, which can cause slightly more neurons to be active). Without this adaptive threshold, a silent attractor dominates retrieval.
2. The notebook then uses the retrieved pattern from (1) to seed a second round of pattern completion using a fixed threshold *θ* = −0.15, which in combination with the global inhibition parameter *γ* = 0.6 provides good retrieval alongside the possibility of retrieving a silent state (see [11] for detailed derivation of performance as a function of these parameters). This two step process enables retrieval of patterns that are not forced to have a fixed sparseness, and a “silent state” attractor can be retrieved when the seeding pattern lies far away from any of the encoded patterns.

This models a simple form of replay. Supposing that the notebook pattern at convergence is 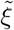, the student input and target output are then reconstructed based on the Hebbian connectivity as 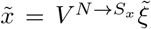 and 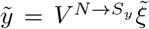. This provides an 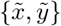 sample from which the student can learn using gradient descent.

The policy is memory-optimized, in the sense that this replay continues indefinitely, such that all samples stored in the notebook are eventually learned by the student. This section characterizes the memory and generalization performance of the student resulting from this replay process. If reactivations perfectly reconstructed the stored examples, this replay strategy would be similar to ‘batch’ learning strategies in machine learning, in which the same stored dataset is repeatedly revisited to update network weights. However, errors in reactivation could in principle degrade the learning process. In Section 6.1 we show that although reactivations introduce errors, remarkably, these errors are correlated in such a way that learning still proceeds like batch learning from perfectly recalled examples up to a rescaling of the learning rate. Using this fact, in Section 6.2 we provide the expected memory and generalization errors, based on results known in prior work [21, 4].

In this policy, both the notebook and student learn potentially beneficial information, and in principle either could be used to answer a specific query for a point x. We take the normative assumption that the best system is selected to make the prediction. Often, this means that the output for a previously stored input will be predicted by the notebook, while that for a novel input will be predicted by the student. However, in Section 6.1 we show that there are conditions under which the student memory error in fact surpasses the notebook, and the student would be used to make predictions for previously stored inputs.

#### 6.1 Accurate learning despite errors in reactivation

How do reactivation errors influence learning dynamics in the student? One hint that learning from reactivations can be effective comes from Fig. 2 of the main text. Given that the notebook is specifically designed to rapidly store memories, it often has a lower memory error than the student. Surprisingly, however, Figs. 2a-h of the main paper show that the student’s training error can fall below that of the notebook. How could it be that the student learns to accurately produce a memory that was imperfectly memorized by the notebook? Our key theoretical observation is that although the notebook imperfectly activates the output of the student, it also imperfectly activates the inputs of the student. These errors are correlated between input and output neurons in a way that does not harm student learning. We demonstrate this fact in this section.

Reactivations have subtly different statistics to the original samples. In particular, when the notebook settles on a pattern *ξ^μ^* (one column of the matrix *ξ*) that was associated with an original sample *x^μ^, y^μ^* from the teacher, this results in reactivated student activity input and output patterns 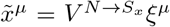 and 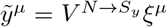, respectively. Horizontally concatenating the input and output reactivations into the matrices 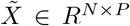 and 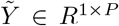, this reactivation leads the weights in the student network to change (in the reactivated gradient direction) by the amount,

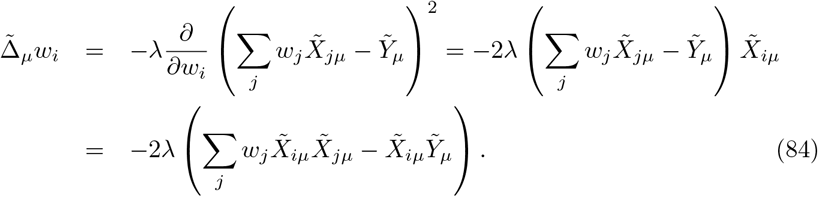

Therefore, the change expected from gradient descent learning with a random notebook index is

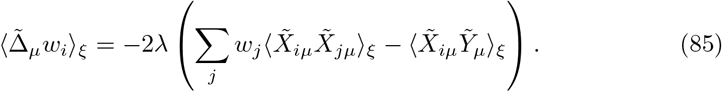

To evaluate these expectations, we form the matrix 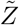 by vertically stacking 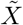 and 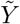, then note that

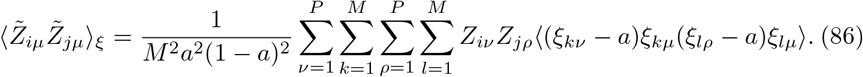

When *μ* ≠ *v* ≠ *ρ*, the statistical independence of memories allows us to factor out 〈*ξ_kv_* – *a*〉, which is zero and causes the whole term to vanish. Similarly, we get no contributions if *μ* ≠ *ρ* = *v*. This implies that both the *v* and *ρ* indices must either pair with each other or with *μ*, and the only terms that contribute are thus *v* = *ρ* = *μ* and *v* = *ρ* ≠ *μ*.

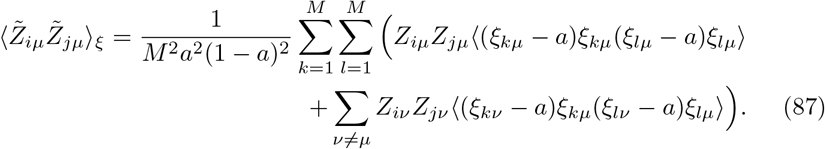

Both of these expectations have been calculated en route to calculating the notebook’s training error. Plugging Eqs. (68) and (72) into the above expression, we find,

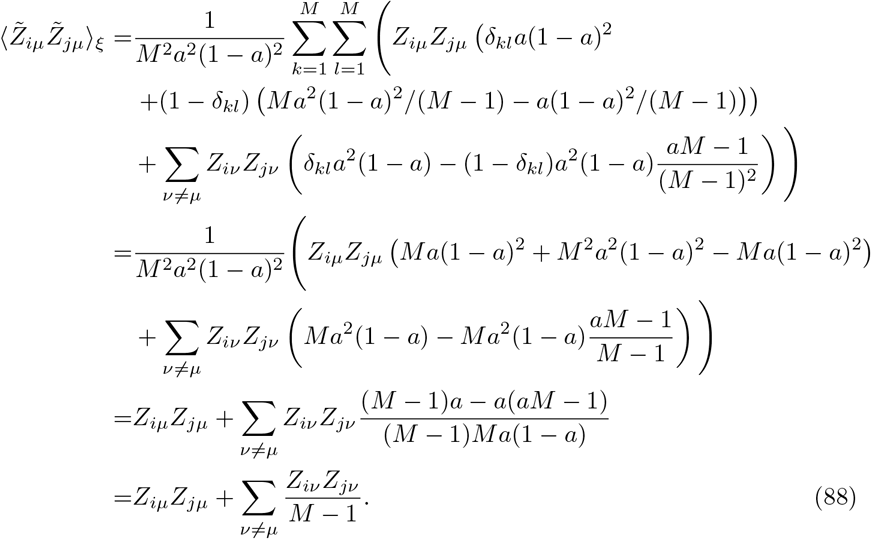

Therefore,

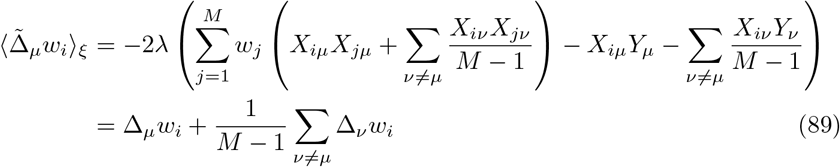

where Δ_*μ*_ *w_i_* is the weight update that would occur if the student were perfectly reactivated by the notebook pattern *μ*. Equivalently, Δ_*μ*_*w_i_* is the weight update that would occur from online learning to the teacher’s example. Importantly, all contributions to 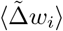 are in the gradient direction of one of the teacher examples. Rearranging this expression slightly, we find:

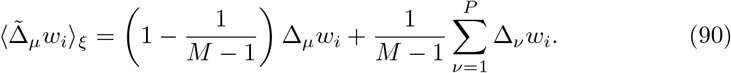

Therefore, each notebook reactivation of pattern *μ* is equivalent to a mini-batch update for that particular pattern with effective learning rate 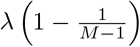, plus a batch update for all stored patterns with effective learning rate 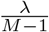. Similarly, the learning expected by sequential notebook reactivation of all *P* patterns is

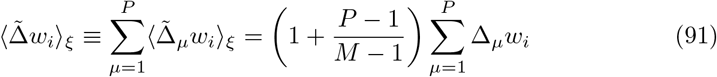

This is equivalent to batch learning with an effective learning rate of

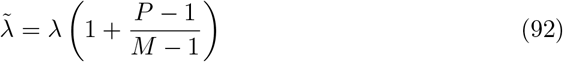

In sum, the notebook’s imperfect reactivation patterns hurt notebook memory performance, but they do not harm the student’s ability to learn from past memories if the learning rate is appropriately controlled.

#### 6.2 Student memory and generalization error from replay

As shown in Sections 5.2 and 6.1, notebook reactivations closely recapitulate stored student activity patterns when run below a critical capacity, and reactivation errors are correlated in such a way as to preserve the relevant statistics for student learning. In this regime, when replay events are random and the learning rate is small, the student effectively learns from the whole batch of samples. Batch learning dynamics differ fundamentally from online learning dynamics, because in the batch setting the noise associated with each example is repeatedly revisited. This difference raises the danger of overfitting to the specific batch of stored data, rather than learning the general rule.

We therefore leverage known solutions to the batch learning dynamics of student-teacher models in our high-dimensional setting [21, 4]. The average memory error is (see Section 2 of [4])

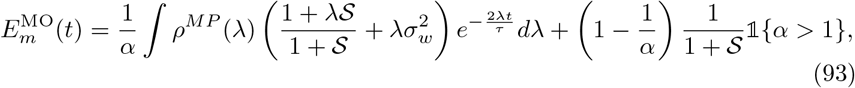

and the generalization error is

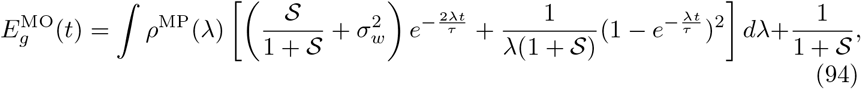

where the superscript MO stands for “memory-optimized,” *t* here measures time in units of epochs, such that each stored example will be replayed once as *t* goes from 0 to 1, 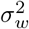 denotes the initialization variance of the student weights, i.e., 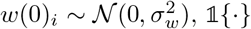 is an indicator function that is 1 when the argument is true and zero otherwise, and the density *ρ*^MP^(·) denotes the Marchenko-Pastur distribution [22, 25], which describes the eigenvalue distribution of the input correlations *XX^T^* in the high-dimensional regime. It has the form

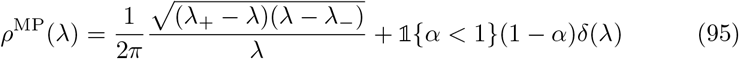

for λ = 0 or λ ∈ [λ___, λ_+_], and is zero elsewhere. The distribution comprises a delta function spike at zero, corresponding to zero-variance input directions that occur when there are fewer samples than the input dimension (i.e., *α* < 1), and a bulk with upper and lower limits, 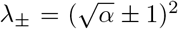, that depend on the load a. We set 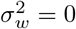 for most analyses in the paper.

We call this strategy memory-optimized, because Eqn. (93) is strictly decreasing in time, so to optimize student memory, replay should be continued indefinitely. However, Eqn. (94) is non-monotonic. Thus, while sustained replay optimizes student memory, this strategy can degrade generalization. Most problematically, it causes catastrophic overfitting at the student capacity, which corresponds to the interpolation threshold where the training error can just reach zero at long times. For a shallow linear student, the capacity is reached when the number of samples is equal to the input dimension, a = 1. Better performance can be obtained for larger and smaller a, a finding known as the *double descent* phenomenon [22, 21, 4, 7]. The behavior of this strategy for a range of SNRs and loads *a* is depicted in Supplementary Fig. S2a,c. While memorization performance is good throughout this space, generalization suffers for low SNRs and loads near one.

#### 6.3 Weight norm dynamics

While memory and generalization error are two key measures of learning progress, we can also ask how the strength of student weights change throughout learning. This quantity could enable certain experimental links, for instance, as a proxy for functional connectivity in the context of the Sweegers et al. [31] experiment discussed in Section 11.

A straightforward modification to the derivation in Section 2.1 of [4] yields the time-dependent average student weight norm as

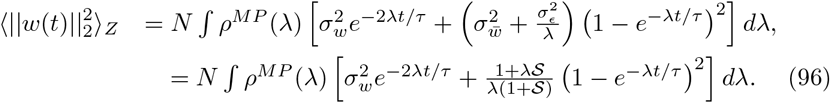

**Figure S2:**
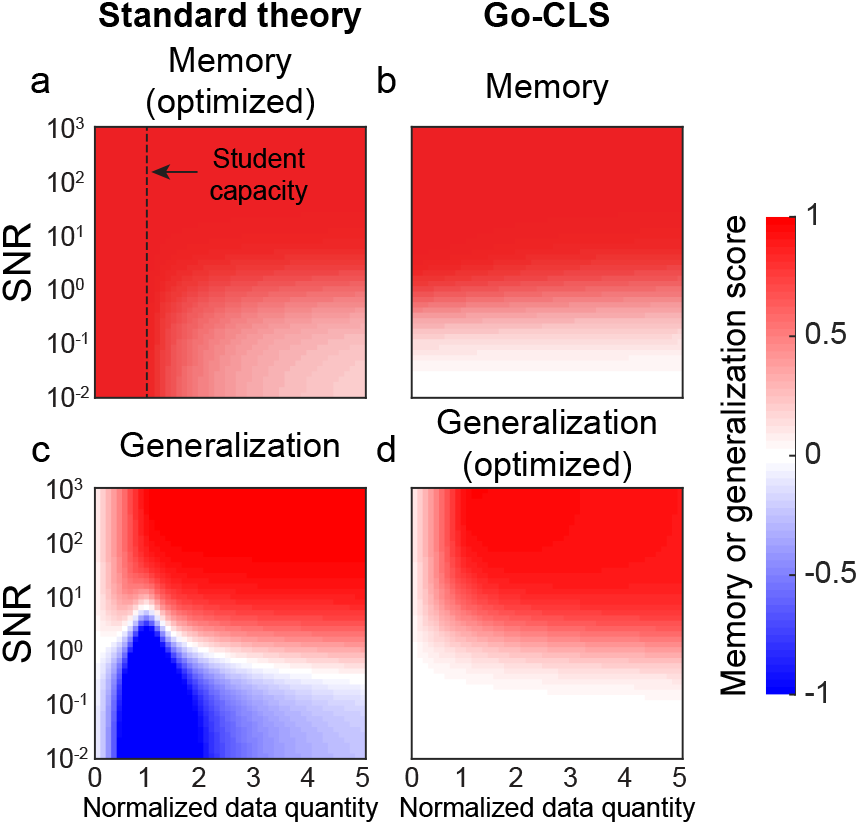
Heatmaps of student memorization performance (a, b) and generalization performance (c, d) as a function of SNR and *α*, when optimized for student memorization (a, c) or generalization (b, d).

Although we typically consider the case where 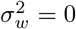, for large 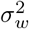, this equation can describe an initial decrease in norm, followed by an increase in norm as weights align with the teacher.

#### 6.4 Correlated training data and non-uniform memory reactivation

Here we numerically explore the effects of introducing input correlations and biased notebook sampling on the training and generalization error dynamics of the student.

In Section 6.1, the errors in notebook reactivation were caused by readout interference when retrieving the training patterns. This effectively introduced correlations in the training data set, and we were curious how correlations affected training dynamics more generally. When 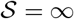, increasing levels of correlation mainly made the generalization error decay slower (Fig. S3a). When 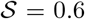, correlated data caused more severe overfitting (Fig. S3b). Interestingly, while introducing correlations in training data generally increased the severity of overfitting, it did not the change our finding that the worst overfitting occurs when the normalized data quantity equals one (*α* = 1) (Fig. S3c). These dynamics could potentially be studied analytically by replacing the Marchenko-Pastur distribution with the eigenvalue distribution of a random matrix ensemble containing uniform input correlations [37].

In Section 6.1, we also assumed that memories were reactivated with uniform probability. This is a good model for the notebook we implemented (Fig. S3d, inset, top), but realistic memory-reactivation mechanisms might be more biased. We simulated non-uniform sampling of memories and found that biased sampling slowed down the rate of generalization error improvement on noiseless data with *P* = *N* (Fig. S3d). On the other hand, it had a small effect in slowing down overfitting in the presence of noise (Fig. S3e). Similar to introducing correlations, biased sampling also did not change where worst overfitting occurred as a function of normalized data quantity (Fig. S3f). We observed subtle changes in the generalization performance under biased sampling compared to uniform sampling, as the trend of the change (i.e. increased or decreased rate of generalization error performance) switched sign as a function of normalized data quantity.

**Figure S3:**
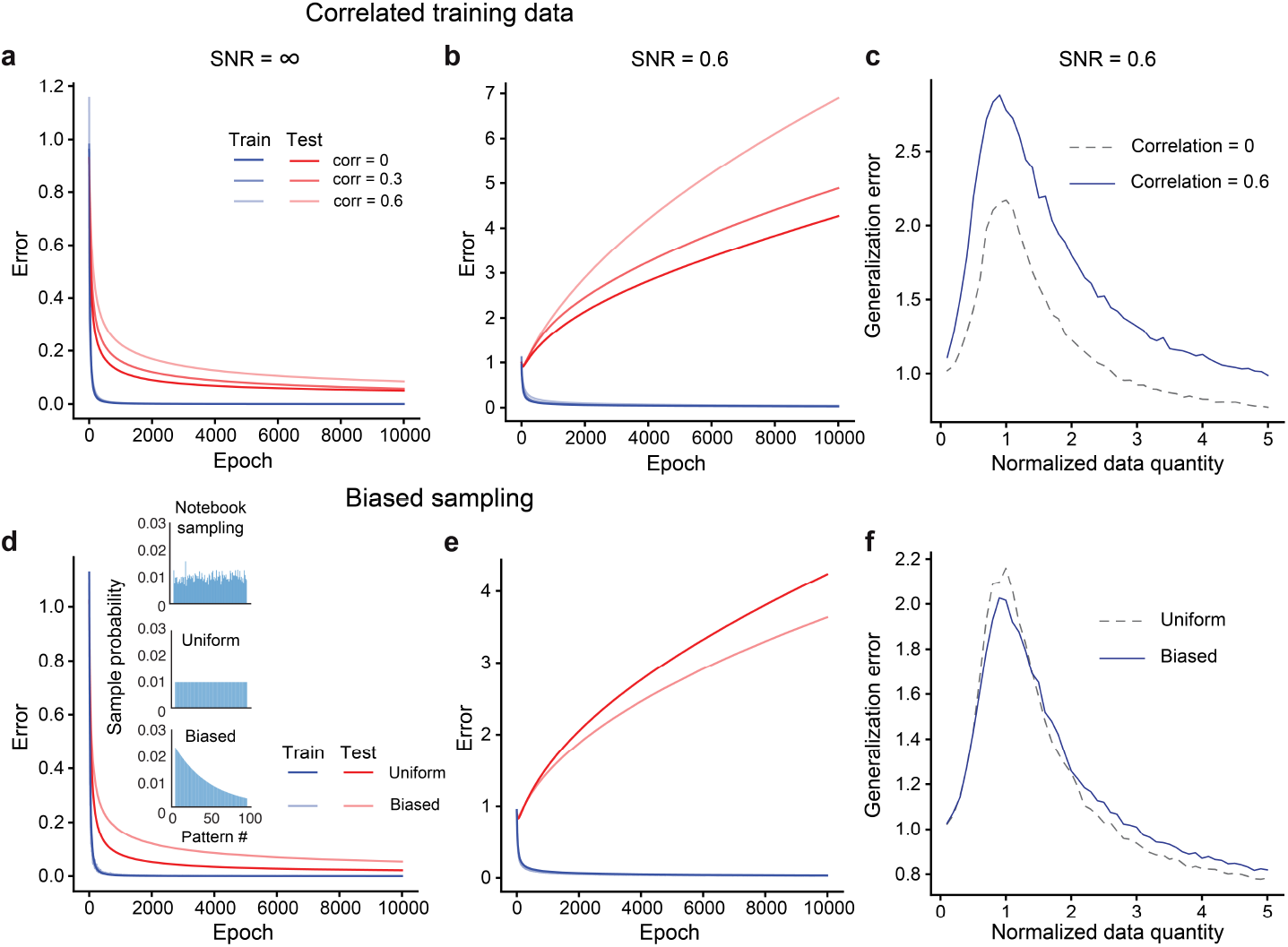
(a,b) Effect of introducing different levels of correlations among the training patterns on the training and generalization error dynamics for 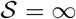 and 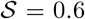, respectively. (c) Generalization error as a function of *α* for both i.i.d training data and correlated training data (generalization error was measured at epoch 2000). (d,e) Effect of biased sampling on training and generalization error dynamics for 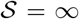 and 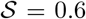, respectively. Biased sampling probabilities were chosen using an exponential function. Inset in d shows the sampling probabilities for training patterns when using the notebook’s random reactivations (one example shown), uniformly sampling, and biased sampling. (f) Generalization error as a function of *α* for both uniformly sampled and biased sampled data. In all simulations, *P* = *N* = 100, and traces are the average of 50 independent runs.

These results extend our main conclusion (i.e. overfitting is a problem when learning from a moderate amount of noisy data) to biased reactivation schemes and alternate noisy data ensembles. They additionally revealed that the severity of overfitting generally depends on more than the SNR and quantity of data, with more severe overfitting here appearing when the training data contained uniform input correlations. Future theoretical analyses could provide a fuller understanding on what other factors influence the dynamics and final results of systems consolidation.

### 7 Generalization-optimized Replay Policy

The generalization-optimized replay policy is similar to the memory-optimized replay policy. Samples are stored in the notebook and replayed to the student to drive learning. When it comes time to make a prediction, the system with the best error is used. The key difference, however, is that replay is not continued indefinitely. Instead, replay is terminated when generalization error stops improving and starts to worsen due to overfitting. That is, this strategy regulates replay to maximize generalization error.

#### 7.1 Student memory and generalization error with regulated replay

In detail, this strategy continues replay until the optimal early stopping time *t*\*, defined as

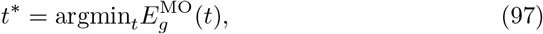

where 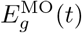 is the memorization-optimized generalization error trajectory (i.e. unregulated trajectory) from Eqn. (94). The student memory and general-ization error therefore have the piece-wise form

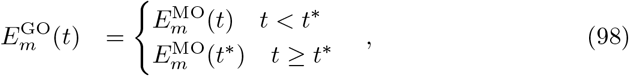

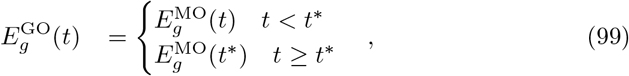

where the superscript GO stands for “generalization-optimized,” and 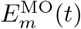 denotes the memorization-optimized memory error trajectory from Eqn. (93). Crucially, under this regulated strategy, the student memory error can remain large indefinitely. Conversely, regulation avoids potentially catastrophic overfitting. The performance of the student under this strategy is depicted in Fig. S2b,d as a function of SNR 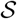 and load *α*. Finally, we note that weight norm dynamics have a similar piece-wise form for this strategy, such that the dynamics follow Eqn. 96 for *t* < *t*\*, at which point they stop.

#### 7.2 Properties of early stopping

To gain a better understanding of this strategy, we can ask how the optimal stopping time depends on dataset parameters. While there is no closed form expression for *t*\*, some intuition can be obtained by computing the optimal stopping time for one fixed value of λ in the integral of Eqn. (94) (a strategy that would be exact if the MP distribution were a delta function at a single value of λ). The optimal stopping time is then (see Sec 2.2 of [4])

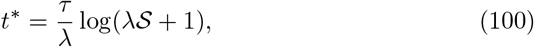

which shows that replay can continue longer for higher SNR relationships, though the relationship is logarithmic.

Early stopping is only one out of a variety of regularization strategies that can combat overfitting. Another possibility is to explicitly penalize large weight values. The *L*_2_ regularization strategy sets the student weights according to

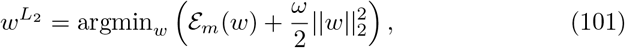

where *ω* denotes the regularization strength. The optimal *L*_2_ regularization strength for our setting is known to be inversely proportional to SNR, 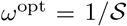 (see [2, 3, 4]). Further, for the specific teacher and student regression problem we consider here, this regularization is known to be Bayes optimal, such that no algorithm can outperform it [2, 3]. It therefore can serve as a normative standard of comparison for early stopping. Prior work has shown that, in our setting, early stopping closely approximates the effect of explicit *L*_2_ regularization (see, e.g., Fig. 5a of [4]), providing a normative basis for the early stopping strategy.

Finally, we can exploit the similar performance of early stopping and optimal *L*_2_ regularization to obtain an explicit (but approximate) expression for the performance of the generalization-optimized replay strategy after the early stopping time. In particular, for *t* > *t*\* we have

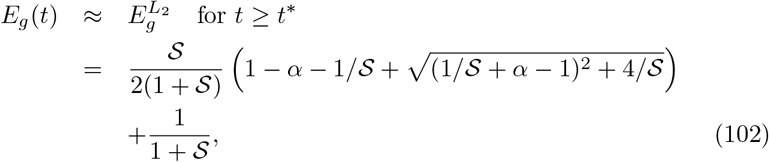

where the latter step is the known generalization error of optimal *L*_2_ regularization on this problem [2, 3, 4].

Using a similar approach, we can approximate the weight norm at the optimal stopping time as the weight norm of the optimal *L*_2_ regularized solution (see Eqn. 66 [4]),

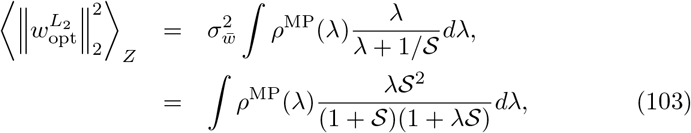

which we note limits to 1 as 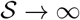 and 0 as 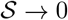, such that high-SNR relationships have larger weight norms than low-SNR relationships at the optimal stopping time.

### 8 Example of generalization non-limiting unpredictability

The main text provides several examples of generalization-limiting unpredictability, with the canonical example being a teacher with output noise. However, not all sources of unpredictability are generalization limiting. For example, suppose that the teacher generates noiseless data,

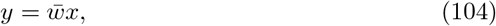

but the student has internal noise in its input neurons that affects its predictions

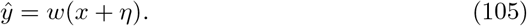

Averaging over the input and noise distributions (but not the teacher weights), the generalization error of the student is

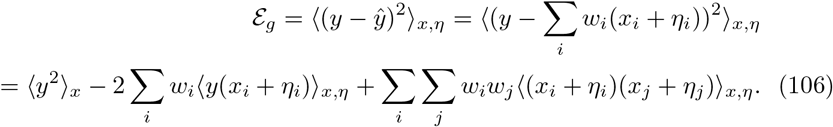

Assuming that *x, y*, and *η* are zero mean random variables, and that *η* is uncorrelated with *x* and *y*, this is equal to

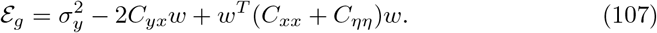

Setting the derivative with respect to *w* equal to zero,

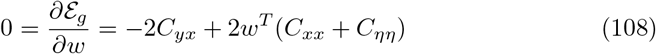

we find that the student weights that optimize generalization are

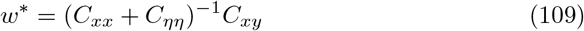

In contrast, the teacher weights satisfy

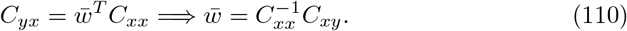

Since 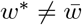, the generalization-optimized student is statistically biased,

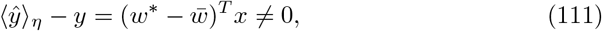

and the generalization error is nonzero. The teacher is unpredictable by the student.

Nevertheless, this type of unpredictability does not require strongly regulated systems consolidation. For example, suppose that the notebook perfectly memorizes *P* input-output patterns of the student. Then, the memory error averaged over student neuron noise is

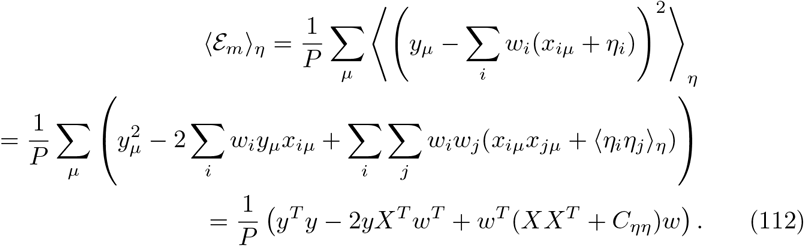

Setting the derivative with respect to w equal to 0,

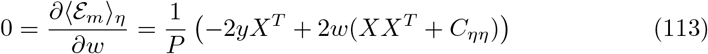

we find that the weights minimizing the training error are

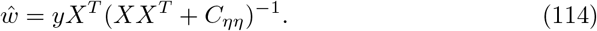

Noting that *XX^T^* and *yX^T^* are (proportional to) estimates of *C_xx_* and *C_xy_* given the *P* teacher examples, we see that this is the same basic form as the weights that minimize the generalization error.

In terms of the learning dynamics, the role played by eigenvalues of *XX^T^* is now played by eigenvalues of *XX^T^* + *C_w_*, which are lower bounded by the minimum eigenvalue of *C_ηη_*. For white noise, this is just 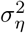. Overfitting was previously due to eigenvalues near 0, but those have now been shifted up to 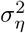. The student input noise regularizes the learning process.

### 9 Means to regulate systems consolidation

Section 7 explained how generalization performance could be optimized by using the SNR of the teacher to regulate the amount of systems consolidation. In biology, the SNR is not known *a priori,* and the brain must decide for itself how to regulate consolidation. This section explores several plausible strategies that the brain could use to regulate systems consolidation. We emphasize strategies that serve to optimize generalization in the teacher-student-notebook framework.

#### 9.1 Validation set approach

In the teacher-student-notebook framework, the notebook stores *P* teacher-generated examples at *t* = 0. We’ve so far assumed that the notebook reactivates all *P* examples with uniform probability to drive student learning. However, it’s also possible for the notebook to divide its examples into separate training and validation sets. As before, the notebook could reactivate the training examples to drive student learning. However, if the validation examples are not reactivated to drive student learning, then they could instead be used to approximate the generalization error. Alternatively, a validation set can be used to train a separate, smaller student. A recent machine learning study shows that such a separately-trained model can be used to construct a score for ranking individual training sample’s usefulness in improving generalization [26], which in turn can be used for regulating consolidation.

#### 9.2 Maximum likelihood estimation

The above strategy used separate subsets of examples to drive learning and estimate the generalization error. Such a scheme could allow the brain the regulate systems consolidation by stopping student learning as soon as the generalization error begins to increase. An alternate strategy is to estimate the SNR of the teacher, 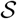, from the examples it provides, *X* and *y*. In this subsection, we calculate and characterize the maximum likelihood estimator,

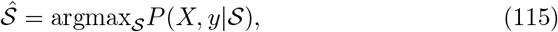

which is a statistically principled and asymptotically optimal unbiased estimator of 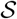. It will be convenient to replace the likelihood function, 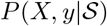, with the log-likelihood function, log 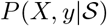, because the logarithm does not change the location of maximum,

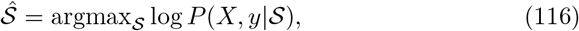

and it is often mathematically convenient to work with log-transformed functions. For example, by the product rule of probability, we have

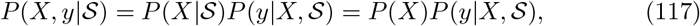

where we noted that *P*(*X*) is independent of 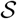 by assumption. Therefore,

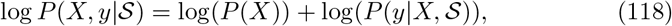

and

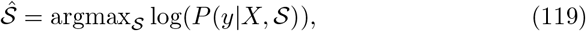

where we discarded 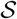 additive factors in the last step.

We next derive an expression for 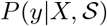. It is convenient to first note that 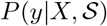 is the marginal of 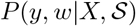 over *w*:

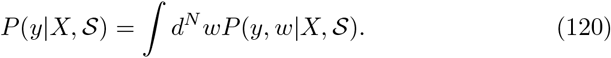

Again using the product rule of probability, we find

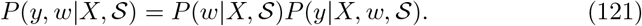

Both of these probability distributions are easy to specify. First, elements of *w* are i.i.d. distributed as 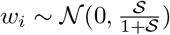 by assumption, so

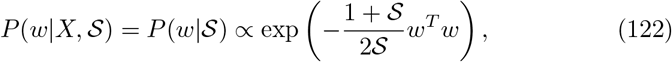

where the normalization constant was neglected for mathematical conciseness and will be put back in later. Second, note that elements of *y* are normally distributed with mean *Xw* and variance determined by the noise, which is i.i.d. distributed as 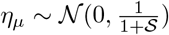. Therefore,

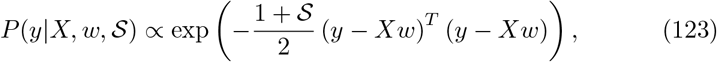

where the neglected normalization constant will again be included later. Putting these two pieces together, we can easily see that 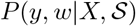 is a zero-mean multivariate Gaussian distribution with an 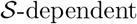 covariance structure. In particular, the arguments of the exponential factors in Eqs. (122) and (123) combine to give

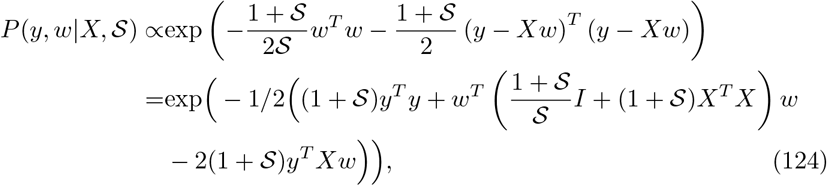

which shows that each term in the exponential is second-order in *y* and *w*. This allows us to use the general Gaussian integral formula to integrate over *w*,

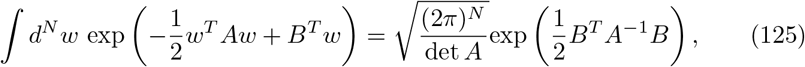

with

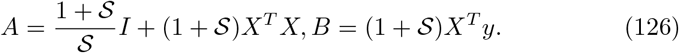

Consequently, 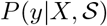 is again a multivariate normal distribution, this time with exponential factor given by

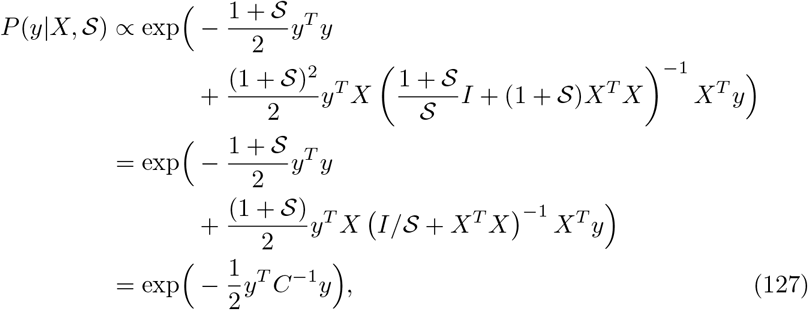

where

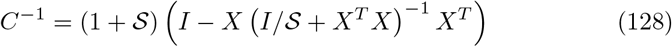

is the inverse covariance matrix. Putting back in the normalization factors, we find

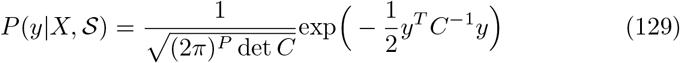

and thus

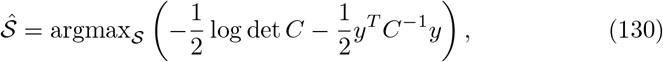

where *C* is a function of 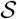 and *X*.

It is convenient and conceptually clarifying to rewrite the covariance matrix in Eq. (128) in terms of the singular value decomposition of *X*,

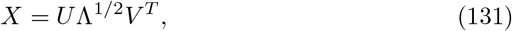

where *U* and *V* are orthogonal matrices and Λ is non-negative rectangular diagonal matrix. In particular, it implies that

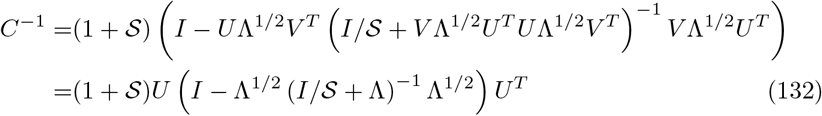

and

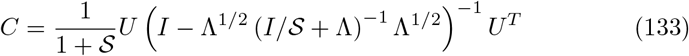

Using the fact that Λ_*ab*_ = λ_*a*_δ_*ab*_ is a rectangular diagonal matrix, this expression can be significantly simplified by recognizing that

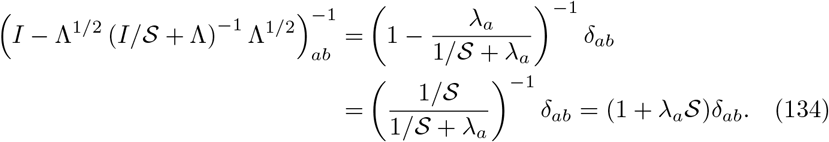

In particular, we see that

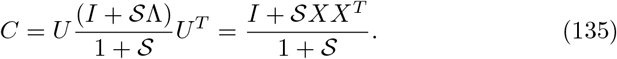

#### 9.3 Learning speed approach

Finally, we consider a simple heuristic based on the initial rate of improvement in a task. Intuitively, tasks with easy-to-memorize data points are those with an underlying pattern that supports generalization (i.e. are high SNR). To formalize this intuition in our setting, we consider the initial slope of the student memory error in Eq. (93),

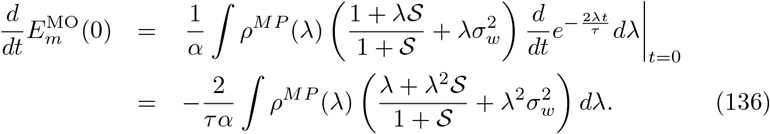

For a given *α, τ*, and 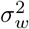, this expression shows that the initial slope depends on the SNR S. Therefore, replay can be optimally regulated by measuring the initial slope (a quantity immediately available to the agent), and reading off the associated SNR. The bottom panel of Fig. 2m illustrates this relationship for one set of parameters, which is approximately linear in the logarithm of the SNR. This heuristic is representative more broadly of a class of approaches that monitor training trajectories as a way of estimating generalization performance.

### 10 Complex teachers

#### 10.1 Linear Student

Here we show that a mismatch between the teacher and student, such that the teacher is deterministic but more complex than the student, is a form of generalization-limiting unpredictability that behaves similarly to observing a teacher with noise. Our derivation follows Appendix C of [4], but we include a derivation of this important result for completeness.

Suppose the teacher generates inputs independently from some distribution *x* ~ *p*(*x*), and labels them using the possibly nonlinear function *y* = *g*(*x*). The best possible linear student (i.e., the student trained on infinite data) will have weights

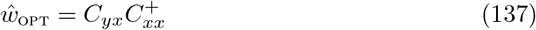

where *C_yx_* = 〈*yx^T^*〉_*x,y*_ is the input-output correlation matrix, *C_xx_* = 〈*xx^T^*〉_*x*_ is the input correlation matrix, and 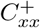 denotes its pseudoinverse. We can rewrite the teacher output as the prediction of this optimal student and a residual,

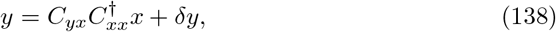

where the residual, *δy*, is defined by this expression.

Next, we consider learning the student weights from a finite batch of data with *P* examples, given in matrices *Y, X* with examples in the columns. The student weights that minimize the training error are

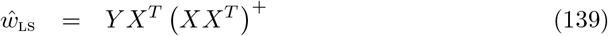

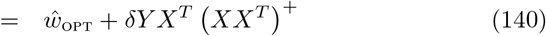

where we substituted Eq. 138 for *Y*, and 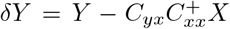 is the matrix of residuals. This formulation clearly separates contributions to the student weights into an optimal component and an overfitting component. Notably, the overfitting *term δYX^T^* (*XX^T^*)^†^ has the same form as for additive noise. This noise is generally non-Gaussian, but the training and generalization errors equal those of a Gaussian teacher with mean and variance matched to *δY*.

#### 10.2 Nonlinear student

In order to test if our theoretical conclusions generalize beyond the linear student setting in regression tasks, we extended our numerical experiments to nonlinear student networks solving real-world classification tasks. More specifically, we trained deep convolutional neural networks (CNNs) to classify images from the MNIST [23], CIFAR-10 [20], and Tiny ImageNet [14] data sets. We used a simple CNN for the MNIST dataset (2 {convolutional + pooling} layers followed by 3 fully connected layers) and ResNet-18 [17] for the CIFAR-10 and Tiny ImageNet datasets (see Fig. S4 associated code for simulation details). For CIFAR-10 and Tiny ImageNet training, data augmentations including random crop, rotation, and horizontal flip were introduced during training. Unaltered examples were used for evaluating training and testing performances.

Interestingly, we found that Go-CLS theory’s essential conclusion still holds for these nonlinear teacher and student cases (Fig. S4). In particular, overtrained networks could achieve full memorization of the training data (100% training accuracy), but these models showed overfitting (Fig. S4a-c), reflected by the increased test loss after reaching a minimum. Similar to the linear student network, early stopping prevented overfitting and resulted in a non-zero training loss (Fig. S4d-f). This reiterates the main message of Go-CLS theory: neural networks trained for perfect memory performance suffer in generalization, and generalization can be improved by regulating the consolidation process according to some regularization scheme (such as early stopping, dropout, or weight decay).

**Figure S4:**
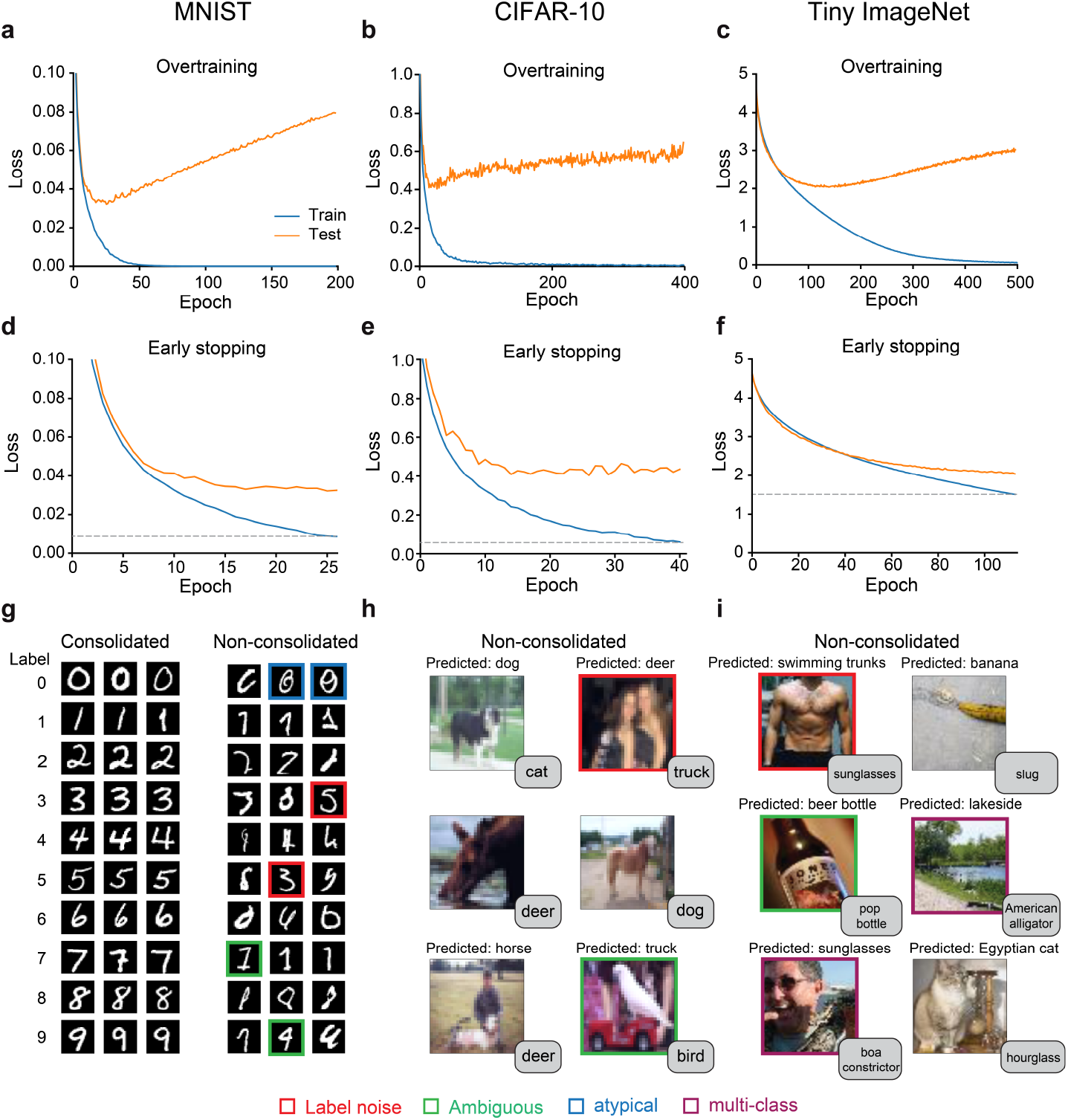
Deep Convolutional Neural Networks (CNN) trained on MNIST, CIFAR-10, and Tiny ImageNet show qualitative similar behavior of overfitting and memorization-generalization trade-off, compared to simpler linear models. (a-c) Deep CNNs can overfit to training data in all three datasets. (d-f) Overfitting can be prevented by early stopping. Gray dashed lines mark the non-zero training loss when performing early stopping. (g) Example training examples that are corrected classified (consolidated) (for g only) versus incorrectly classified (non-consolidated) after early stopping. For h and i, the text box at the bottom right corner of the image shows the ground truth label.

A novel aspect of these real-world data sets is that individual examples can have different noise characteristics. Examining the training examples that were correctly classified (consolidated) versus incorrectly classified due to early stopping (non-consolidated) provides interesting hints about the nature of the data that can harm generalization if fully consolidated. Taking the MNIST task as an example, the consolidated digits are easily recognizable as ‘canonical’ examples of each class (Fig. S4g, left), with variations that typically did not cause ambiguity (e.g. 7 vs 7 with a slash through it). By contrast, non-consolidated digits exhibited several distinct forms of unpredictability. First, there are clearly mislabeled data in the MNIST dataset (digits with red boxes). These digits act as noise during training, mirroring the “noisy teacher” discussed in the main paper (Fig. 5b). Second, many digits seemed to be written in ambiguous ways or atypical ways (e.g. examples of 0 with a slash through it). This suggests that more information is sometimes needed to determine the right label (i.e. about the writer’s penmanship style), making these examples akin to the “partially observable teacher” (Fig. 5d). Similarly mislabeled and ambiguous data are also abundant for the unconsolidated images in CIFAR-10 and Tiny ImageNet datasets (S4h, i). These data sets additionally contain non-consolidated images that contain multiple distinct objects within the same scene. For example, one image contains both a cat and a hourglass, but the network’s classification of “Egyptian cat” is counted as incorrect.

### 11 Comparison of Go-CLS theory to past experimental results

Go-CLS theory generates a diversity of amnesia curves that might help memory researchers explain the similar diversity found experimentally (Fig. S5). Researchers usually classify hippocampal amnesia dynamics according to whether memory deficits are similar for recent and remote memories (flat retrograde amnesia), more pronounced for recent memories (graded retrograde amnesia), or absent for both recent and remote memories (no retrograde amnesia) (Fig. S5a). Since real world experiences are composed of many elements that differ in their degree of predictability (Fig. 5j), our theory predicts that different components of human memory will consolidate to different degrees (Fig. 5k). In human memory research, patients with selective hippocampal damage indeed show retrograde amnesia reflecting diverse dynamics of systems consolidation [38, 16]. Some patients show graded retrograde amnesia consistent with the standard theory, while others either have flat retrograde amnesia or no retrograde amnesia [38] (Fig. S5b). Similarly diverse retrograde amnesia curves have been seen in rodent memory tasks (Fig. S5c-f). For example, hippocampal lesions can result in either graded or flat retrograde amnesia in different individuals performing the same task [35, 30, 36] (Fig. S5c-e), and individual animals can exhibit different types of amnesia on different tasks [36] (Fig. S5e,f).

Go-CLS theory recasts this wide range of experimental observations through the tuning of two parameters (Fig. 3e), the predictability of experience and the amount of prior consolidation. It is not yet possible to unambiguously specify these parameters for arbitrary real-world experiences and experimental memory tasks, but the empirical patterns are plausibly consistent with Go-CLS theory. For example, famous faces and facts about public events are generally reliable components of many life experiences, and one need not conjure up a specific past experience to remember what Barack Obama looks like or that the COVID-19 pandemic stunned the global economy. Memories of famous faces and facts about public events may thus represent content that is highly predictable across experiences, in which case Go-CLS would predict that they can be consolidated. Indeed, many patients can recall remote facts and famous faces without a functioning hippocampus [38] (Fig. S5b). In contrast, autobiographical memories combine idiosyncratic details about specific experiences in one’s life that may not generalize to other experiences. For example, one often needs to think back to the original experience to remember the cake served at their child’s birthday party or the songs played at their wedding. Because many incidental influences shape how complex real-life events unfold, remembering autobiographical memories may require the recall of content that is intrinsically unpredictable. Go-CLS predicts that unpredictable content will not be consolidated, and most patients cannot recall these memories without a hippocampus [38] (Fig. S5b). Along similar lines, the Morris water maze task consistently requires the hippocampus [36, 10, 28] (Fig. S5f), perhaps suggesting that rodents need to recall past experiences to reconstruct the detailed arrangement of environmental cues and platform positions [27, 29], both chosen arbitrarily by the experimenter. Rigorously assessing these *post hoc* interpretations will require theoretical and experimental progress on the algorithms used for predictability estimation.

Go-CLS theory can also generate diverse time courses for time-dependent generalization that mimic experimental diversity [35, 34, 13, 8]. For example, some mice showed increased fear responses to similar but not identical contexts in fear-conditioning experiments (“generalizers”, Fig. S5g, red bars), while others maintained distinct behavioral responses over time (“discriminators”, Fig. S3g, blue bars) [35]. Strikingly, only the discriminators required their hippocampus for memory recall of the original context (Fig. S5h). Although other interpretations are possible, it’s intriguing that Go-CLS theory predicts that memory transfer and generalization improvement should be similarly correlated (Fig. 3f, S5i, S5j). In this interpretation, “discriminators” might judge fear conditioning to be an unpredictable experience that should not consolidate because this would cause maladaptive generalization. Their memories would thus be left in original form and be susceptible to strong retrograde amnesia (Fig. S5i, j, blue bars). In contrast, “generalizers” might infer that the experience is predictable, which would then lead to consolidation, weak retrograde amnesia, and learned generalizations (Fig. S5, red bars). This variability across individuals is possibly due to differences in each animal’s regulation process (Fig. 2i-m) or feature encoding (Fig. 5i).

**Figure S5:**
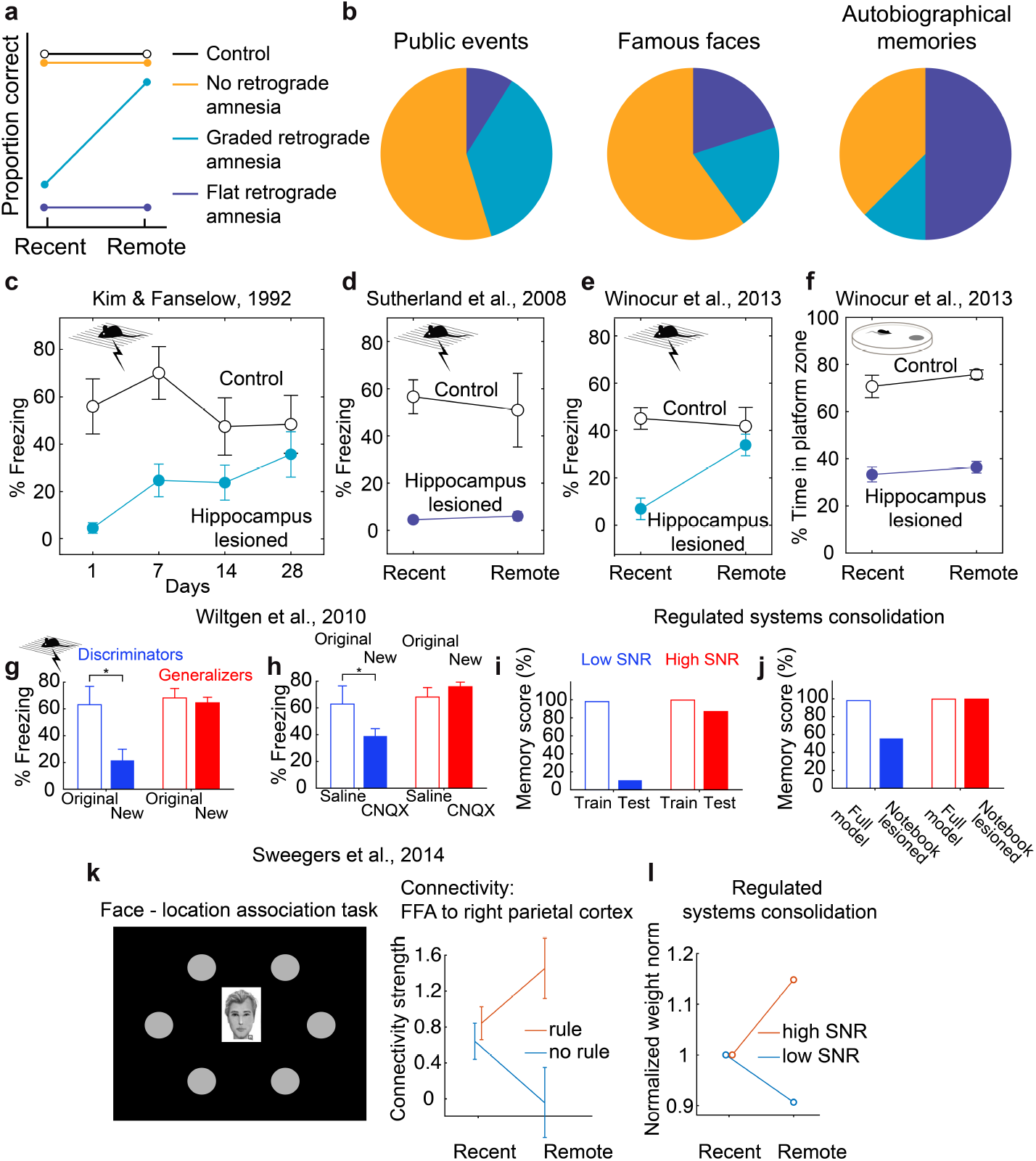
Diverse findings in memory research. (a) Schematic of retrograde amnesia curves. (b) Reports of retrograde amnesia in human patients with selective hippocampal damage show diverse dynamics. Figure adapted from Yonelinas et al., 2019 [38]. (c, d) Lesioning hippocampus in rodents can produce both graded and flat retrograde amnesia. Figure adapted from Kim & Fanselow, 1992 [19], Sutherland et al., 2008 [30]. Lesioning the hippocampus can result in graded (e) or flat (f) retrograde amnesia in the same animal performing different tasks (contextual fear conditioning and Morris water maze, respectively). Figure adapted from Winocur et al., 2013 [36]. (g) Discriminators can differentiate the original fear-conditioning context with another similar but novel context, whereas generalizers show similar amount of fear response to both contexts. (h) Silencing the hippocampus in mice 15 days after contextual fear conditioning differentially impact fear memory of the original context, depending on whether the animal show time-dependent fear generalization. panels g and h are adapted from Wiltgen et al., 2010 [35]. (i, j) Regulated systems consolidation can reproduce similar correlation between time-dependent generalization and reduced hippocampal dependence of memories. High SNR (1000) and low SNR (0.6) simulations based on analytical solutions are used to model the “generalizers” and “discriminator”. 2000 total epochs are simulated with N = P = 100, notebook size M = 5000, and learnrate = 0.005. (k) Face-location association task with rules vs no rules show different time-dependent change in functional connectivity between cortical areas. Figure adapted from Sweegers et al., 2014 [31]. (l) Regulated systems consolidation shows similar connectivity changes over time, as reflected in the norm of the student’s weights. Student weight w is drawn i.i.d. from N(0, 0.5), where the weights’ non-zero initial condition reflect the brain’s preexisting connectivity between these two regions. The student then learns from a high SNR teacher (SNR = 2) or a low SNR teacher (SNR = 0.05), while the weight norm is monitored through time (normalized to the initial norm). Note that a decrease in weight norm is expected on the low-SNR learning task, as a large weight norm generates substantial output variance that is uncorrelated with the teacher’s noisy output. 2000 total epochs are simulated with N = P = 100, notebook size M = 2000, and learnrate = 0.015.

An experiment closely related to Go-CLS theory was performed by Sweegers et al. [31]. In their task design, healthy human participants had to associate specific faces with positions on a computer screen (Fig. S5k, left). Half of the locations were assigned faces through an unpredictable random process, whereas the other locations were assigned faces according to a hidden but fully reliable rule. The authors then used functional magnetic resonance imaging (fMRI) to assess how systems consolidation changed the functional connectivity between several brain areas during memory recall. They specifically asked whether functional connectivity patterns revealed statistical interactions between the association type (faces assigned to rule locations versus no-rule locations) and time (recall at recent versus remote time points). We subsequently refer to these statistical interactions as rule/time interactions.

Sweegers et al. found selective cortical recruitment that is consistent with the general premise of Go-CLS theory. More specifically, they detected statistically significant rule/time interactions in the functional connectivity from the hippocampus to a region containing parts of the anterior cingulate cortex (ACC) and medial prefrontal cortex (mPFC) [31]. This suggests that the time course of system consolidation depends on the predictability of the consolidated information. A *post hoc* analysis revealed similar rule/time interactions in the functional connectivity from this ACC/mPFC region to the fusiform face area (FFA). These findings collectively indicate enhanced ACC/mPFC connectivity for the rule locations at the remote time point, consistent with the idea that systems consolidation is regulated to recruit neocortical computations when the predictability of experience is high.

Intriguingly, Sweegers et al. also described a trend in their data that could quantitatively link their experimental paradigm to Go-CLS theory. In particular, when they investigated how functional connectivity from the fusiform face area (FFA) differed at recent and remote time points, they found that its connectivity to right parietal cortex increased for the rule-based locations and decreased for the no-rule locations (Fig. S5k, right) [31]. This result is expected from Go-CLS theory (Fig. S5l). The right parietal cortex is involved in spatial processing, and we interpret its functional connectivity from FFA in the teacher-student-notebook model as student weights used to predict neural activity coding location from neural activity coding faces. Our theory predicts that the predictability of the face-location relationship determines whether systems consolidation drives neocortical learning that links FFA to right parietal cortex. Indeed, these connections strengthened only when the face-location relationship was predictable. This empirical difference can be quantitatively captured by regulated systems consolidation (Fig. S5l). However, the authors did not test the statistical significance of this trend, because earlier analyses failed to detect any rule/time interactions in the functional connectivity from FFA. As such, the authors were worried that subsequent statistical tests would have inaccurately inflated p-values. It would be very interesting to see whether this trend is significant in a replication of Sweegers et al.

### 12 Experimentally testable prediction of Go-CLS

In this section, we provide some predictions of the Go-CLS theory and outline a framework for designing experiments to test these predictions. The core feature of these testable predictions is that subjects performing any task should consolidate information that is highly predictable, but not information that is unpredictable. Thus, any experimental design requires a way to vary the predictability of an input-output task and to measure the amount of consolidation that occurs during learning. Having accomplished this, it should be possible to design experiments that probe the mechanistic implementation of regulated consolidation, provided that the initial experiments are indeed consistent with Go-CLS. In theory, experiments can be done on any species capable of learning input-output tasks with variably predictable relationships. Here, we focus on mammalian species with hippocampal and neocortical brain regions (e.g., humans, non-human primates, rodents, etc.), as consolidation is presumed to occur through interactions between these two brain regions. Below, we provide a general recipe for designing such tasks. We presume that the details of experimental design will be determined by domain experts who wish to perform these types of experiments.

#### 12.1 Testing the relationship between predictability and consolidation

The essential component is a behavioral task consisting of an input-output relationship designed by the experimenter and learned by the subject through experience. To design such a task, the experimenter must first choose inputs (e.g., visual cues), outputs (e.g., sound frequencies), an action to indicate the output (e.g., movement to target), a relationship between the inputs and outputs that can take predictable and unpredictable forms, and a reward (or removal of aversive stimulus) that serves as the driver for the subject to learn the input-output relationship. Predictable here means that if the subject sees either an old or a novel cue, it can use a rule learned from past trials to make a good prediction about how to respond correctly. The unpredictable version of the task lacks a systematic input-output relationship, and the only strategy for good performance is to memorize which cues are rewarded and which ones are not. Thus, we refer to the predictable and unpredictable versions of the task as *rule-based* and *memory-based* tasks, respectively. In addition to these two versions of the task, levels of generalization performance should be quantifiable by testing the performance on novel experiences following the same generative process. This is a critical piece of information on whether the animal is indeed consolidating predictable information in the manner we have originally designed.

Furthermore, there must be ways to infer the extent of systems consolidation. A key method is to determine if performance is impaired by hippocampal dysfunction (preferably reversibly) at remote time points. Our theory predicts the extent of memory preservation after hippocampus dysfunction should be correlated with the predictability of experience. Manipulations such as lesions and reversible silencing (chemical, optogenetic, chemogenetic) can be applied to experimental animals (e.g., rodents) at different time points post-learning. This is more difficult in humans because of the constraints associated with using invasive or genetic tools for hippocampal perturbation, but it might be possible with functional imaging methods. For example, following systems consolidation of rule-based (highly predictable) experience (i.e., in a well-trained subject), memory recall is predicted to not require the hippocampus, and memory recall may thus engage the hippocampus more weakly. Strong neocortical activation must occur while the subject performs the task, and we expect higher within-neocortex functional connectivity. In contrast, weaker within-neocortex functional connectivity would be expected during performance of a memory-based (unpredictable) version of the task than the rule-based version, which relies more on reactivation of hippocampal memories to make predictions. Alternatively, perturbations of the hippocampus could be potentially achieved in humans through fMRI-guided transcranial magnetic stimulation (TMS) [32].

In the following, we provide two example experiments following the above-prescribed recipe:

Example 1: Visual gratings presented at various angles are used as the inputs and colored reward ports are used as outputs. Animals can indicate their choice by licking at a particular reward port (Fig. S6a). A relationship with varying degrees of predictability can be introduced to link the input and output. For example, the high predictability version of the relationship could be that angles from 0° to 45° correspond to licking at the blue reward port, angles from 45° to 90° correspond to licking at the yellow reward port (Fig. S6a). For a task with low predictability, the mapping between angles and reward ports could be selected randomly (i.e., no systematically predictable relationship between angle and reward port, but the mapping could be memorized) (Fig. S6b). Our theory predicts that, over time, the hippocampus is no longer required to solve the rule-based (highly predictable) version of the task due to full consolidation (i.e., hippocampal silencing will not impair the correct performance of either the learned examples or the novel examples following the same rule). On the other hand, in the task with randomly established mapping, silencing the hippocampus would impair the animal’s ability to recall previously memorized grating-to-reward-port mappings. Note that specific choices of such input/output pairs and their mapping rules would require considerations of the species-specific biology and ethology. For example, humans and non-human primates might be able to perform the angle-based task well, whereas mice might not have the visual acuity to differentiate nearby angles [1].
Example 2: An experiment performed by Sweegers et al. (Fig. S6c) is consistent with our task design recipe. Specifically, the inputs are human faces with certain features, the output is one of six locations on the screen to which the experimental subject needs to move a joystick to indicate the choice. The mapping was designed to be either highly predictable, mapping certain features to certain locations, or unpredictable, consisting of a random mapping between features and locations. The authors in this study measured systems consolidation using the strength of cortico-cortical coupling for remotely acquired associations compared to recent ones, indicated by functional connectivity (see Supplementary Material 11). fMRI-guided TMS could potentially be used in future studies to perturb the hippocampus for a more causal investigation of the relationship between task predictability and hippocampus dependence.

**Figure S6:**
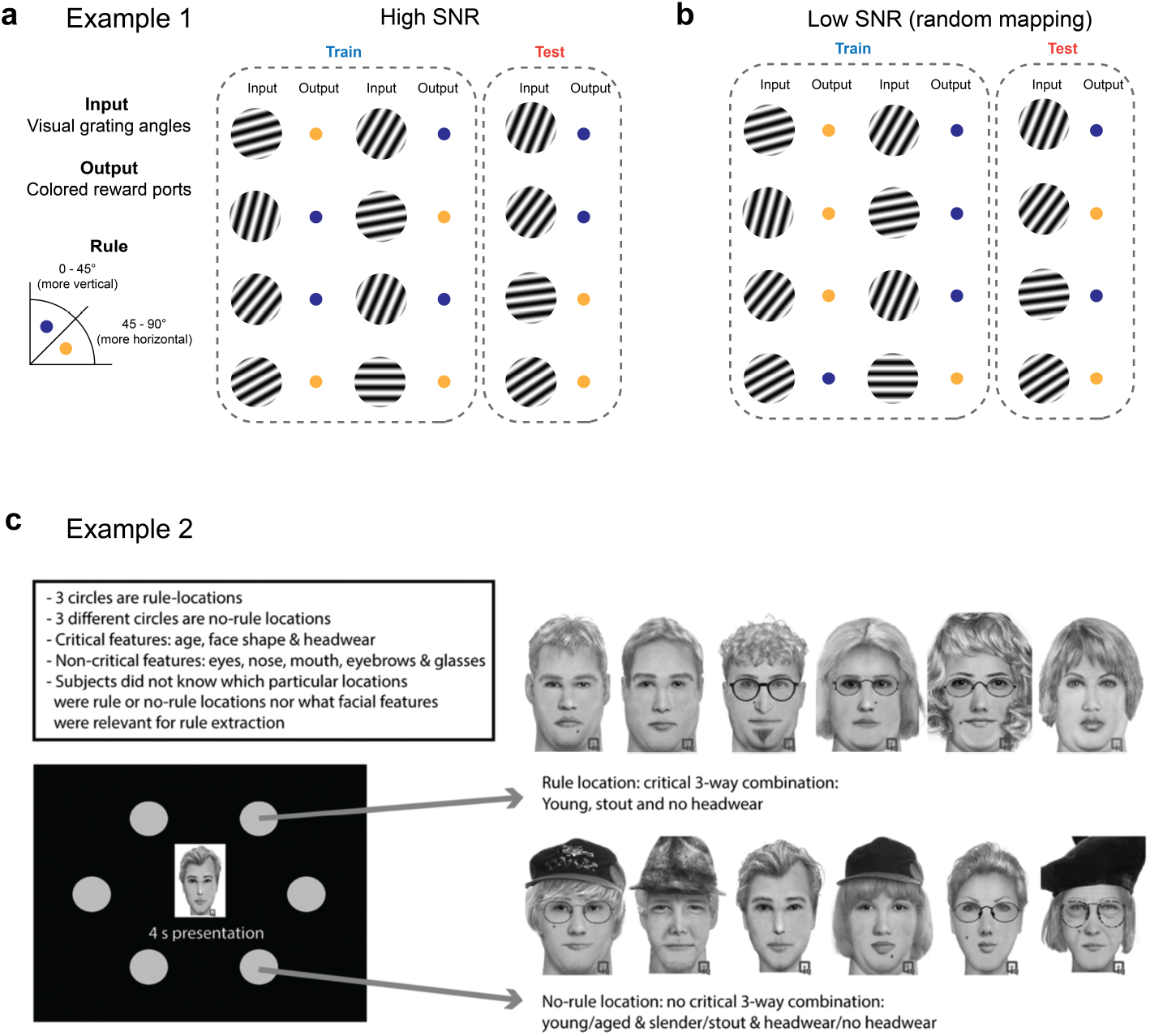
Example experiments for testing Go-CLS. (a, b) Illustrations of a rule-based and a memory-based input-output task. (c) Task design for a facelocation task used in Sweegers et al., 2014 [31].

#### 12.2 Experiments probing implementational mechanisms of Go-CLS

If experiments like the ones described above yield results that are consistent with the Go-CLS theory (i.e., show clear correlation between the level of systems consolidation and the task predictability), implementational mechanisms for how systems consolidation is regulated can then be studied. Experimental tools for investigating possible mechanisms will differ by species. In humans, non-invasive tools such as fMRI, electroencephalography (EEG) or magnetoencephalography (MEG) will be the predominate methods for monitoring brain activity. In animal model systems, finer-scale measurements can be performed in combination with the behavioral experiments described in the previous section. For example, electrical recordings using electrode arrays (e.g., tetrode arrays or neural pixel probes) or cellular-resolution calcium imaging can be used to record neural activity. The tools available in any of these species permit experiments to monitor changes in the way representations of the task change over time during learning in the hippocampus, neocortex, or both. Spontaneous neural activity during resting periods can also be recorded in any of these species to decode task-related representations [24] and monitor correlated changes in offline activities and the extent of consolidation. For model systems like rodents, optogenetic or chemogenetic tools can be used to causally identify different brain regions involved, perhaps even prior to recording/imaging, in order to focus those efforts on the most pertinent brain regions. Manipulations can be timed to interfere with learning during training sessions or during offline periods when consolidation is expected to occur. Similar experiments in humans could potentially be done through fMRI guided TMS or deep brain stimulation.

Once critical brain regions and appropriate manipulation times are identified, further refinement of experimental protocols can be used to dig deeper into mechanisms. For example, hippocampal sharp wave ripples (SWRs) associate with offline replay of prior experiences and have been shown to have a causal role in mediating systems consolidation [12]. It would therefore be natural to monitor these replay events (e.g., through electrophysiology in rodents or MEG or fMRI in humans [24]) after the subject learns either the rule-based or memory-based tasks. Replay content can then be decoded to see if rule-based experiences are replayed more than memory-based experiences. If so, this would be consistent with replay regulation as a mechanism for content-dependent regulation of systems consolidation.

If rule-based experiences are indeed preferentially replayed, it would then be possible to study the underlying mechanisms that selectively bias these replay events. One possibility is that brain regions outside of the hippocampus send inputs to the hippocampus to bias the content of replay, based on predictability. For example, in mice, the medial prefrontal cortex (mPFC) can influence hippocampal activity indirectly through the thalamic nucleus reuniens. mPFC has been implicated as a crucial brain region in rule-learning and cognitive control, and it is plausible that mPFC contains the predictability information needed to bias replay content. Two predictions follow from this hypothesis. First, neural activity in mPFC would be fundamentally different during offline replay of rule-based versus memory-based experiences. Second, silencing thalamic nucleus reuniens or mPFC (e.g., by chemogenetics) would disrupt the regulation of replay in a manner dependent on the predictability of experience. Other brain regions (e.g., ventral tegmental area) could be studied in a similar manner. Once such brain regions are identified in rodents, more detailed mechanistic experiments can be performed to map the cellular level connectivity, cell types, neurotransmitters, receptors, and other molecular processes involved in the regulation of systems consolidation.

On the other hand, it is also possible that hippocampal replay content and frequency are similar for experiences with differing predictability. In this case, an alternative hypothesis would be that regulation of systems consolidation occurs outside of the hippocampus. For example, replay events could be regulated so that neocortical regions respond differently depending on predictability of the content. This hypothesis is theoretically appealing because memories are composed of many relationships that differ in their predictability. To test this hypothesis, experimental techniques such as wide-field or mesoscopic calcium imaging (rodents) or fMRI (humans) can be used to monitor cortical-wide activity during periods of offline replay. Differential activation of the neocortical regions between the rule-based and the memory-based tasks would be informative about whether systems consolidation is regulated at the neocortical level.

## Footnotes

1 To avoid confusion, recall that *η^αN^* notates the learning rate for example *αN* ≈ *μ*, and this notation is not meant to imply an exponential learning rate schedule.

2 Note that this argument neglected the threshold present in the notebook dynamics. This choice reflects the fact that we want the argument to carry over to the linear student neurons that we will consider in the next section. Nevertheless, pseudoinverse weights also work for thresholded notebook neurons in the typical case that *f* (−*ξ*) = 0 and *f* (1 – *θ*) = 1.

